# Event-triggered MINFLUX microscopy: smart microscopy to catch and follow rare events

**DOI:** 10.1101/2025.08.27.672674

**Authors:** Jonatan Alvelid, Agnes Koerfer, Christian Eggeling

**Affiliations:** Institute for Applied Optics and Biophysics, Friedrich Schiller University Jena, Jena, Germany; Leibniz Institute of Photonic Technology, Jena, Germany; Department of Applied Physics and SciLifeLab, KTH Royal Institute of Technology, Stockholm, Sweden; Jena Center for Soft Matter, Jena, Germany

## Abstract

MINFLUX microscopy is a powerful microscopy method allowing for the characterization of molecular organization and dynamics with single nanometer spatial resolution and sub-hundred microseconds temporal resolution. However, acquisition times often span minutes to hours as a single fluorophore is measured at a time. Applying it to study cellular processes in living cells therefore requires careful consideration of where and when to apply MINFLUX data acquisition, a consideration where manual control limits its potential applications. Here, to overcome the limitations of acquisition speed, acquisition initiation, and data throughput, we present a smart microscopy method that uses confocal imaging as a monitoring method, runs real-time image analysis, and only applies MINFLUX data acquisition exactly where and when deemed necessary based on the analysis outcome. The method, event-triggered MINFLUX, is controlled through a custom-written and open source Python widget that automatically controls a commercial MINFLUX microscope. We apply this method to investigate molecular membrane dynamics and organization during three different cellular events: two-dimensional lipid dynamics at caveolae; three-dimensional membrane topography during dynamin-mediated endocytosis; and three-dimensional membrane fluidity and topography during budding site formation of HIV-1 proteins. Thanks to rapid event detection and minimal regions of interest the method provides data that would be unfeasible or impossible to acquire through manual control of the microscope.

## Main

MINFLUX (minimal fluorescence photon fluxes) microscopy^1–4^ excels in spatial resolution and temporal tracking resolution, where it also in commercial instrumentation reaches single nanometers and below 100 microseconds^5^. The power comes at the cost of lengthy recording times of minutes to hours as only a single fluorophore is tracked at a time. Applications have thus so far ranged from imaging bacterial^6^ or synaptic^7^ proteins in fixed cells, to tracking individual motor proteins^8–10^ or cargo transport through the nuclear pore^11^. So far MINFLUX has only sparsely been applied in living cells to capture short-lived seconds-scale events or tracking faster molecules during events of interest, and not for the likes of endocytosis and tracking lipids in cellular membranes.

Smart microscopy^12,13^ and sample-adaptive acquisition approaches has helped conventional and super-resolution microscopy methods to improve light exposure, recording times, image quality, and deep-tissue imaging^14–20^. Multiscale and multimodal approaches have been applied to improve conventional microscopy^21^, STED (stimulated emission depletion) microscopy^22,23^, SMLM (single-molecule localization microscopy)^24^, light-sheet microscopy^25,26^ and SIM (structured illumination microscopy)^27^, and have allowed imaging of both large-scale and small-scale events, slow and fast, from embryo development^28^, through cellular division^29–31^, to synaptic vesicle recycling^22^. Event-triggered methods^14,21,22,27^ are of particular interest, thanks to their ability to spot fleeting biological events otherwise difficult to capture with their respective microscopy methods. As an important aspect, open source software has in many cases been released to allow wide-spread adaptation of the techniques^22,24,29,31,32^. Hitherto, no smart microscopy methods have been applied to MINFLUX, where the potential benefits of increasing the throughput and speeding up acquisitions by limiting regions of interest temporally and spatially are vast^4^. Such a method could enable MINFLUX to achieve a higher throughput of repetitive cellular events, and access rare cellular events on the seconds timescale, providing 2D and 3D data of unrivalled quality.

Here, we present event-triggered MINFLUX (etMINFLUX) microscopy where MINFLUX acquisitions (2D or 3D, imaging or tracking) at the site and time of an event of interest are triggered by the outcome of real-time analysis of confocal measurements. Confocal imaging up to 1 Hz is used to monitor the sample for an event of interest, and when captured the microscope switches to a MINFLUX acquisition. We apply the method to three different biological systems and cellular events, proving the versatility of etMINFLUX: firstly, the throughput of lipid tracking at caveolae sites is highly increased; secondly, endosomes budding from the membrane are detected and mapped in 3D on a timescale of tens of seconds; and thirdly, Gag-protein accumulation sites as an indicator of HIV-1 (human immunodeficiency virus) budding sites are detected and the membrane fluidity and shape is followed over a timescale of minutes.

## Results

### Event-triggered MINFLUX method

The smart microscopy scheme etMINFLUX is capable of triggering 2D and 3D MINFLUX imaging or tracking immediately upon detection of cellular events in confocal images (Fig. 1a). The triggering happens on the scale of hundreds-of-milliseconds, and a region of up to 80 × 80 µm^2^ can be monitored with confocal imaging. The resulting confocal images are analyzed with a pre-determined real-time analysis pipeline specifically optimized to detect the events of interest in the current sample. When an event is detected, such as the accumulation of a fluorescent signal, the confocal imaging is paused and MINFLUX is applied in the immediate surrounding of the event site (Fig. 1b). The size of the region of interest (ROI) can be pre-determined or decided by the analysis pipeline, depending on the type of event being detected. In general, we apply ROI sizes of 0.6–2 × 0.6–2 µm^2^, a crucial detail in MINFLUX acquisitions that e.g. in the application of tracking diffusing molecules allows for rapid sampling of the full ROI on the order of tens of seconds. After the MINFLUX acquisition has finished, after a predetermined or flexible time, the full event data is saved, including the confocal time lapse leading up to the detected event, the MINFLUX data, and a log file containing metadata about the analysis pipeline used, the event detection, and timings. The full metadata and confocal data can be used to validate the event detection in post processing. After the data saving, the microscope automatically returns to the confocal mode and continues the process from the top. etMINFLUX is implemented in a way to allow for indefinite acquisitions without manual interference, and the sample stabilization system of the microscope further ensures data quality throughout.

**Figure 1.**
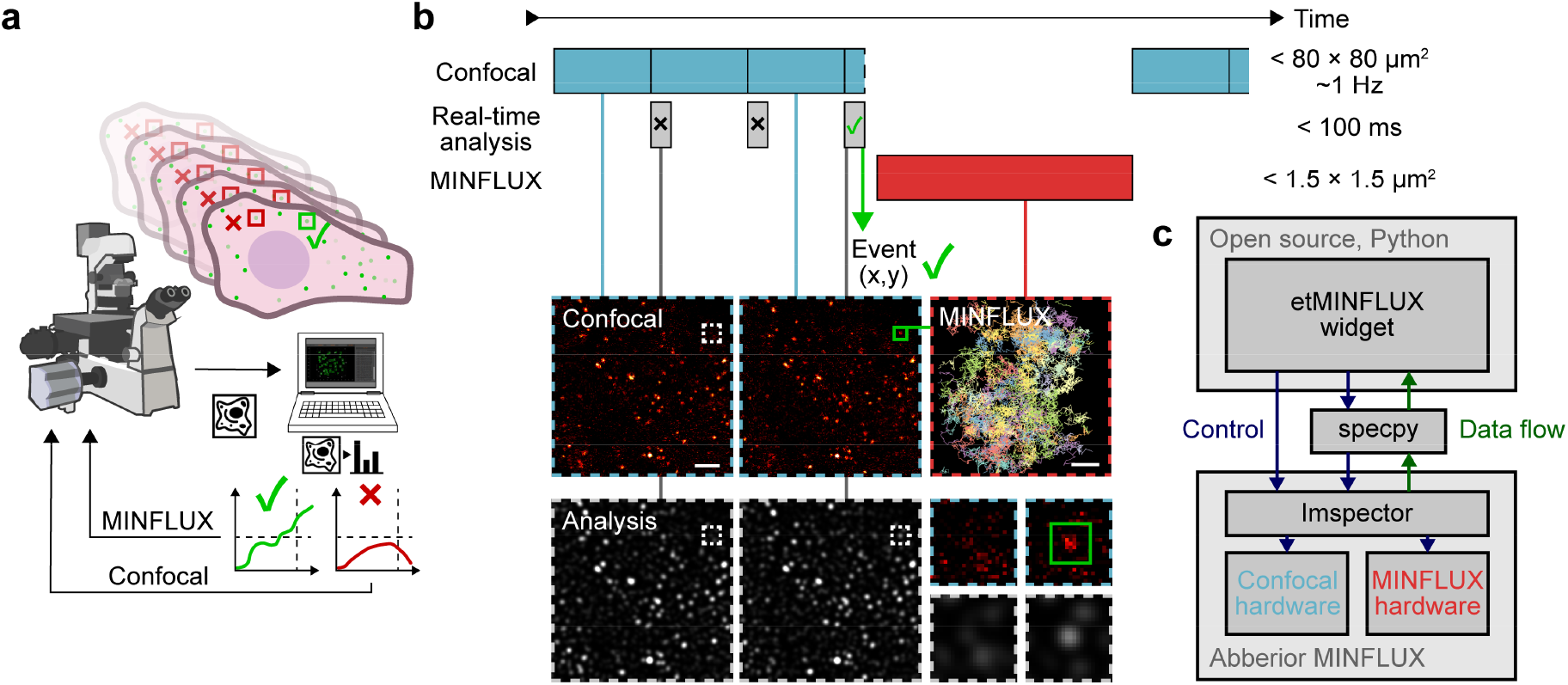
Overview of event-triggered MINFLUX and software widget. **a**. Simplified sketch of the etMINFLUX method. **b**. Timeline sketch and timings and sizes of an etMINFLUX experiment (top), with confocal imaging (top, blue), real-time analysis (middle, gray), and MINFLUX tracking or imaging (bottom, red). Example confocal images, processed analysis images, and a MINFLUX tracking dataset (bottom). **c**. Block diagram of microscope, microscope control software, and etMINFLUX control widget. Scale bars: 2 µm (confocal in **b**), 200 nm (MINFLUX in **b**).

etMINFLUX is implemented as a standalone Python-based and open source control widget, interacting with the Imspector control software of the commercial abberior MINFLUX microscope (Fig. 1c). The control occurs mainly through the *specpy* Python API package for Imspector, while lack of functionality in *specpy* is solved through other direct interactions by the widget with Imspector (Suppl. Note 1). Confocal data is immediately upon acquisition pushed to the control widget, a process controlled using line-update signals. This allows data to be pushed after each line of acquisition and not just full frames, further allowing an increase in event detection speed where necessary. Confocal data is immediately processed through a real-time analysis pipeline, with parameters controllable in the widget GUI (Suppl. Fig. 1). The implementation is flexible to the type of analysis to apply, implementing it as a general Python function and thus allowing the user to flexibly add new event detection types.

For flexibility we developed four different recording modes (Fig. 2, Suppl. Fig. 2). The simplest form of etMINFLUX is a single MINFLUX acquisition following the detection of a single event (Suppl. Fig. 2). In the second mode, multiple event sites are simultaneously identified in real time and MINFLUX acquisitions are subsequently applied at ROIs surrounding each event site for a fixed amount of time (Fig. 2a). The third recording mode deals with following an event site for a prolonged time, where cycles of interleaved MINFLUX and confocal acquisitions are performed at a pre-set confocal frame rate following an event detection (Suppl. Fig. 2). Here, both the monitored signal in the confocal channel and the MINFLUX channel can be tracked as they evolve over acquisition cycles and time. In the fourth recording mode, multiple event sites can be followed simultaneously (Fig. 2b). Here, a time window or threshold for the number of event sites for event detection is determined, and detected event sites are added to a queue until the goal is met. When the goal is met, or the user manually starts the MINFLUX acquisitions, interleaved MINFLUX at the individual event sites and confocal recordings are cycled, as such following the development at all sites simultaneously. Such a recording mode is beneficial to increase the data throughput when the ROI sites should be followed for a longer time. The implementation of recording modes is readily extensible.

**Figure 2.**
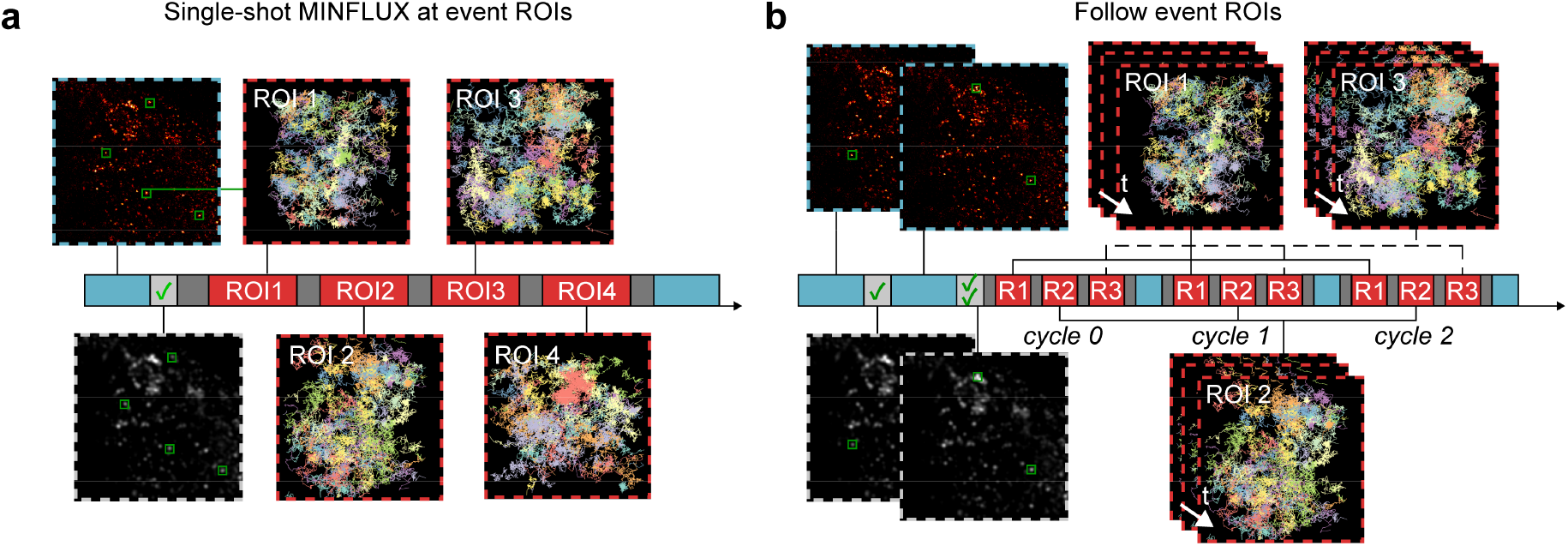
Experimental modalities enabled by event-triggered MINFLUX. **a**. Multiple event ROI recordings after detecting multiple event sites simultaneously. **b**. Multiple event ROI following after detecting multiple events in an event detection time window, with interleaved confocal and MINFLUX recordings of all sites. Timescales show confocal acquisition (blue), MINFLUX acquisition (red), real-time analysis (light gray) and overhead (dark gray). Confocal images are shown with look-up table Hot, and processed images from the analysis pipeline are shown with Gray look-up table Gray. MINFLUX tracks are colored according to their track id.

During any etMINFLUX experiment, the user is free to interact with the acquisition and can apply changes such as moving directly to the next ROI or moving to the next recording cycle (Suppl. Fig. 1). A subwidget further shows the full stack of confocal images leading up to events in a zoomed-in area as well as an intensity graph of the local intensity changes. Another subwidget allows the user to interact with the queue of ROIs that will be recorded, where individual ROIs can be deleted, for example if the event detection was false or if the cellular event has ended.

To finally overlap the confocal and MINFLUX data, for further analysis, a residual relative shift between the coordinate systems of the two methods occasionally has to be applied to ensure trustworthy analysis. The shift depends on the scanning speed of the confocal image, as we can approximate the galvanometric mirrors as stationary during the MINFLUX acquisition, and the relative position in the applied scanning range of the galvanometric mirrors. Characterization and calibration of shift curves depending on scanning speed and position has been performed, and shifts of up to 100 nm are applied to the data where necessary (Suppl. Note 2, Suppl. Fig. 3).

### etMINFLUX to measure lipid diffusion at caveolae

We applied etMINFLUX to measure lipid diffusion at caveolae sites in PtK2 cells with a high throughput (Fig. 3a). Caveolae are abundant plasma membrane invaginations of 60–80 nm in diameter of varying depth, mainly formed by caveolins and cavins^33,34^. They are known to be involved in a range of processes involving signal transduction, endocytosis, mechanoprotection, as well as lipid regulation^35^, but many questions around their role still remains unanswered. While caveolae are dynamic structures, both in formation, disassembly, and scission, as well as movement on the membrane, many are static over a timescale of minutes^36^. Therefore, it served as an initial test for the method, both for the first single ROI and second multiple consecutive ROIs single-shot recording modes (Fig. 3b). Specifically, we expressed fluorescent-protein tagged Caveolin1 (Caveolin1-EGFP) in live cells, and following a confocal image of Caveolin1-EGFP at a size of 15–80 × 15–80 µm^2^, which took 0.3–3.2 s, the *peak_detection* analysis pipeline (Suppl. Note 3) was run, taking on average 69 ms (Suppl. Fig. 4), to detect peaks of the Caveolin1-EGFP fluorescence signal. As caveolae sites were detected, a fast 2D MINFLUX tracking acquisition with a triangle targeted coordinate pattern (TCP) of a dye-lipid conjugate of STAR RED and either sphingomyelin (SM) or 1,2-dipalmitoyl-sn-glycero-3-phosphoethanolamine (DPPE) was performed for around 60 s in a region of 1–2×1–2 µm^2^, before either moving to the next detected caveolae or back to the confocal for a new detection round. Resulting MINFLUX tracking metadata distributions for the temporal track length (62 ± 111 ms), time between localizations (345 ± 888 µs, minimum 86 µs, median 153 µs), and localization brightness (213 ± 102 kHz) for a representative raw dataset are plotted in Suppl. Fig. 5. In total, thanks to the automation, hundreds of ROIs surrounding individual caveolae and random control sites were measured (Fig. 3c). A vast majority (92%) of ROIs showed at least one lipid passing through the site as analyzed with a diameter of 100 nm, indicating the power of limiting the acquisition to a small region to rapidly sample the measured space: acquiring a larger area spanning multiple ROIs would lead to a significantly lower throughput of the data of interest and would increase the risk of the site dynamically moving during MINFLUX acquisition due to longer recording times.

**Figure 3.**
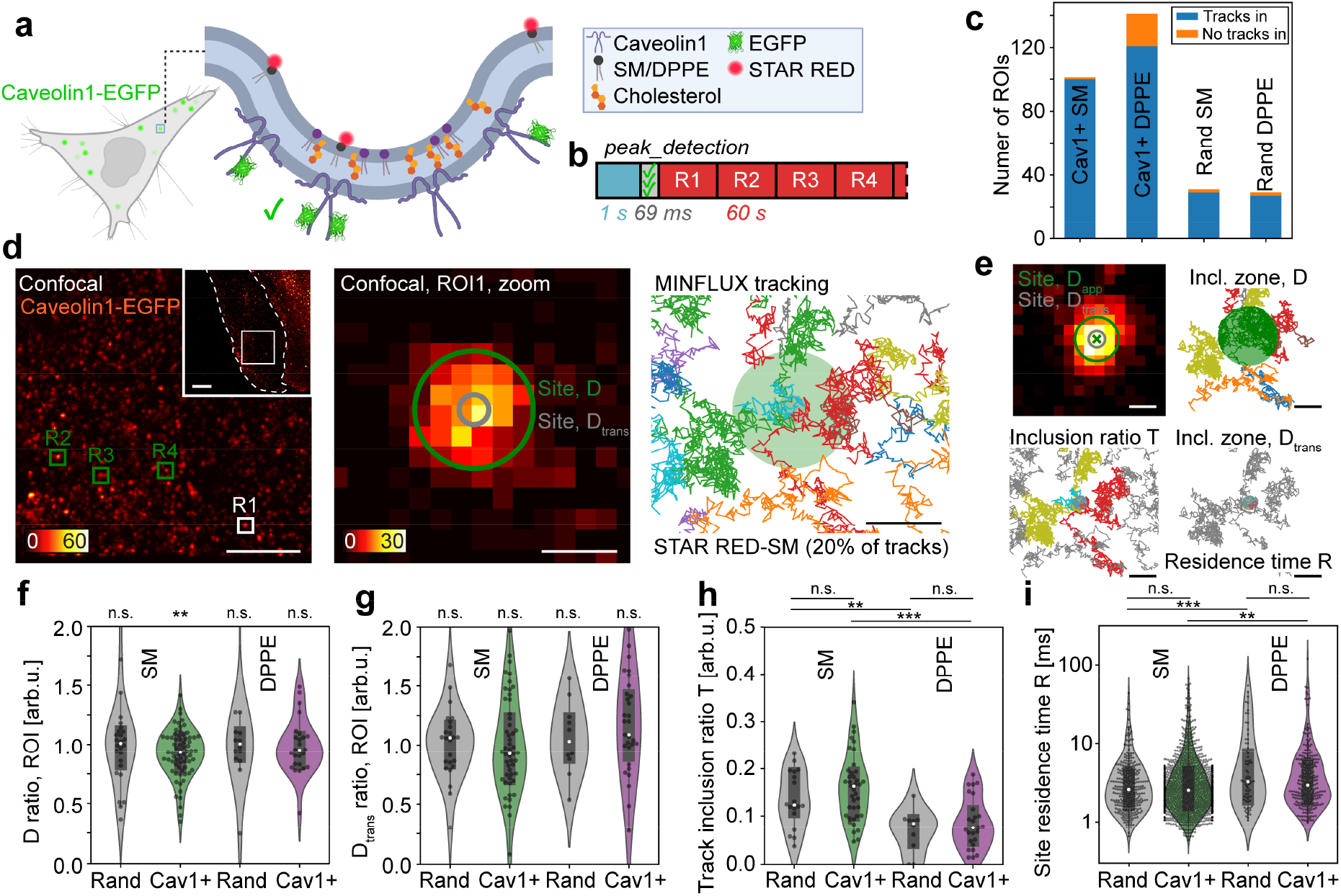
Lipid dynamics measurements at caveolae sites following event detection of Caveolin1 sites. **a**. Sketch of a caveolae site, including experimental Caveolin1-GFP and STAR RED lipid labelling of either sphingomyelin (SM) or DPPE. **b**. etMINFLUX experiment timeline, with confocal (blue), analysis (light gray), and MINFLUX (red) blocks. Indicated times refer to mean (analysis) or median (confocal, MINFLUX) times over all experimental replicas. **c**. Number of analysed ROIs for the four different experimental conditions, including number of ROIs showing tracks inside (blue) or no tracks inside (orange). C+ SM, caveolae sites with labelled SM; C+ DP, caveolae sites with labelled DPPE; R SM, random sites with labelled SM; R DP, random sites with labelled DPPE. **d**. Exemplary experimental data, showing full confocal image of Caveolin1-EGFP (first column), zoom-in to six ROIs selected at caveolae sites (second column), MINFLUX tracking data of STAR RED-SM or STAR RED-DPPE at ROIs (third column), and overlaid MINFLUX tracking and confocal data at ROIs (fourth column). **e**. Visualization of analysis concepts: inclusion zone radii used for the various diffusion analysis (top left); track sections marked as inside (green) and outside (gray) the inclusion zone for diffusion analysis (top right); tracks marked as passing (green) or not passing (gray) the inclusion zone for track inclusion analysis (bottom left); and track sections marked as inside (green) and outside (gray) the inclusion zone for transient diffusion analysis as well as residence time R analysis (bottom right). **f**. Diffusion coefficient ratio analysis, for ratio between diffusion inside and outside the inclusion zone, in the four datasets. Each datapoint represents the mean of one caveolae site. **g**. Transient diffusion coefficient ratio analysis, for ratio between diffusion inside the inclusion zone and outside the inclusion zone for diffusion coefficient analysis, in the four datasets. Each datapoint represents the mean of one caveolae site. **h**. Track inclusion ratio T analysis, for ratio between the tracks that pass the inclusion zone and the total number of tracks, in the four datasets. Each datapoint represents the mean of one caveolae site. **i**. Site residence time R analysis in the four datasets. Each datapoint represents one track situated in a site. **f–i**. Datasets: C+ SM, caveolae sites with labelled SM; C+ DP, caveolae sites with labelled DPPE; R SM, random sites with labelled SM; R DP, random sites with labelled DPPE. Scale bars: 10 µm (overview in **d**), 5 µm (large confocal in **d**), 250 nm (confocal zoom in **d**; MINFLUX tracks in **d**; **e**).

Single or multiple ROIs in the same cell were acquired sequentially after a single confocal detection (Fig. 2a, Suppl. Fig. 2), and the confocal and MINFLUX data was overlaid (Fig. 3d). Following a check that the Caveolin1 accumulation site did not move during MINFLUX recording using a post-MINFLUX confocal image when available, MINFLUX track data filtering steps, and fitting of the precise caveolae site position in the confocal data (Suppl. Note 4, Suppl. Fig. 6), the rich data allowed us to perform lipid diffusion analysis as well as flagging lipid tracks and track sections that were present in the site (Fig. 3e, Suppl. Fig. 6). Firstly, we performed square displacement analysis, and fitting of the resulting datasets to obtain diffusion coefficients for the lipid tracks and parts of those tracks (Suppl. Note 5). From the square displacement fitting analysis, we (in these and similarly all later experiments) further extracted the dynamic localization precision. In this way, the 2D localization precision was determined to be 9.5 nm (Suppl. Fig. 7). Using only the tracks that at some point passed through a marked site (as defined by a circle with a radius of 200 nm; Suppl. Note 5, Suppl. Note 6, Suppl. Fig. 8), we divided these tracks into continuous sections that stay either entirely inside or outside the caveola. For each track section, we performed a square displacement analysis and extracted a diffusion coefficient. Doing this for all tracks at one site, we extracted a mean diffusion coefficient for the sections inside and outside (Suppl. Fig. 9), and calculated a diffusion coefficient ratio for each site as the diffusion coefficient inside divided by the diffusion coefficient outside. We compared the results from random non-caveolae sites, where we expect a ratio of 1, with those from caveolae, for SM and DPPE (Fig. 3f). The ratios at random sites, both for SM and DPPE, were indeed centered at 1 (SM: 1.01 ± 0.07, p = 0.918; DPPE: 1.06 ± 0.12, p = 0.624). The ratios for SM diffusion at caveolae showed a significant decrease of diffusion speed inside the caveolae (0.92 ± 0.02, p = 0.002), while DPPE does not show a difference (0.98 ± 0.04, p = 0.688). This indicates an inhibition of diffusion of SM in caveolae, while DPPE was not significantly affected compared to elsewhere on the membrane.

We further calculated the transient diffusion coefficient for each localization by square displacement analysis and fitting of the resulting distribution in a window (100 localizations) around said localization (Suppl. Note 5). For an identified caveola, we calculated the average diffusion coefficient for localizations inside the site, as segmented with a diameter of 100 nm, and those outside (Suppl. Fig. 9), again extracting a ratio between the two for each site (Fig. 3g). For the transient diffusion analysis, the expected ratio around 1 was observed for random sites with SM (1.01 ± 0.06, p = 0.575) and DPPE (1.06 ± 0.09, p = 0.710), while there was a non-significant trend towards lower ratios for SM (0.95 ± 0.06, p = 0.217) but not DPPE (1.05 ± 0.14, p = 0.620). To further investigate the involvement of the lipids at sites, we analyzed the track inclusion ratio T by taking the ratio between the number of tracks in a ROI that pass through the caveola and the number of tracks that do not pass through (Fig. 3h). The larger the ratio, the more the lipid was in theory enriched in the measured caveola compared to a lipid with a lower ratio. For the selected caveolae sites, we observed a higher inclusion ratio T of SM compared to DPPE (SM: 0.16 ± 0.01, 0.09 ± 0.01; p = 0.00001), i.e. there is an enrichment of SM in caveolae compared to DPPE. The control measurements on random (non-caveolae) sites revealed a decreasing trend of the inclusion ratio compared to the caveolae sites for both lipid analogues (SM, random: 0.14 ± 0.01, SM caveolae: 0.16 ± 0.01, p = 0.110; DPPE, random: 0.07 ± 0.01, DPPE caveolae: 0.09 ± 0.01, p = 0.187), and a higher inclusion ratio of SM compared to DPPE also for the random sites (p = 0.002). This results from the fact that diffusion of SM and DPPE is in general, even outside caveolae, different (Ext. Data Fig. 4–5)^37–39^.

Lastly, we investigated the site residence times R for track sections that passed through the caveola, extracting the time between the first localization upon entry and the last localization upon leaving the site (Fig. 3i). Here, we observed mean R for the 100 nm diameter regions of around 4–8 ms (SM random: 3.84 ± 0.26 ms; SM caveolae: 4.17 ± 0.17 ms; DPPE random: 7.79 ± 1.26 ms; DPPE caveolae 5.61 ± 0.63 ms). DPPE did in general stay longer in caveolae than SM (p = 0.002), and the same tendency was present for random sites (p = 0.000002), which cannot be explained by slower diffusion rates (Suppl. Fig. 9). Most importantly, neither SM (p = 0.355) nor DPPE (p = 0.110) resided significantly longer in caveolae than at random sites.

### etMINFLUX 3D tracking topography of budding endosomes in live cells

In a second biological system, we applied etMINFLUX with 3D MINFLUX tracking of a cholesterol-derived membrane probe, STAR RED-membrane, to measure the topography of the membrane at Dynamin1-accumulation sites in HeLa cells (Fig. 4a). Dynamin is known to accumulate and act as the initiator of scission at clathrin-coated pits, caveolae, and other endocytosis sites^40^, where cholesterol further is important for membrane curvature^41^ and thus abundantly present. The process of accumulation of dynamin and scission is a fast process with respect to MINFLUX acquisitions, occurring on a time scale of tens of seconds^42,43^. Here, we highlight that etMINFLUX was able to catch processes in living cells not feasible to catch with normal MINFLUX microscope control and acquisitions due to their rarity and rapidity – in this case budding endosomes before scission. We performed 3D tracking with an octahedral targeted coordinate pattern (TCP) of the cholesterol-based fluorescent membrane probe STAR RED-membrane, hypothesized to be incorporated in the site due to the local cholesterol-enriched membrane, in order to get a 3D topographic map. Resulting MINFLUX tracking metadata distributions for the temporal track length (46 ± 89 ms), time between localizations (485 ± 626 µs, min 357 µs, median 359 µs), and localization brightness (172 ± 80 kHz) for a representative raw dataset are plotted in Suppl. Fig. 5. We used the single ROI following mode, to ensure the fastest sampling with MINFLUX of a single site (Fig. 4b). Following confocal images of a roughly 15×15 µm^2^ area, with frame rates of around 1 Hz, the *dynamin_rise* real-time analysis pipeline (Suppl. Note 3) was applied, taking on average 26 ms (Suppl. Fig. 4). Confocal imaging was continued until a Dynamin1 accumulation event was detected, at which point the system switched to MINFLUX 3D tracking acquisition of the fluorescent cholesterol-based membrane probe with recording windows of around 20 s in around 0.5×0.5 µm^2^ ROIs.

**Figure 4.**
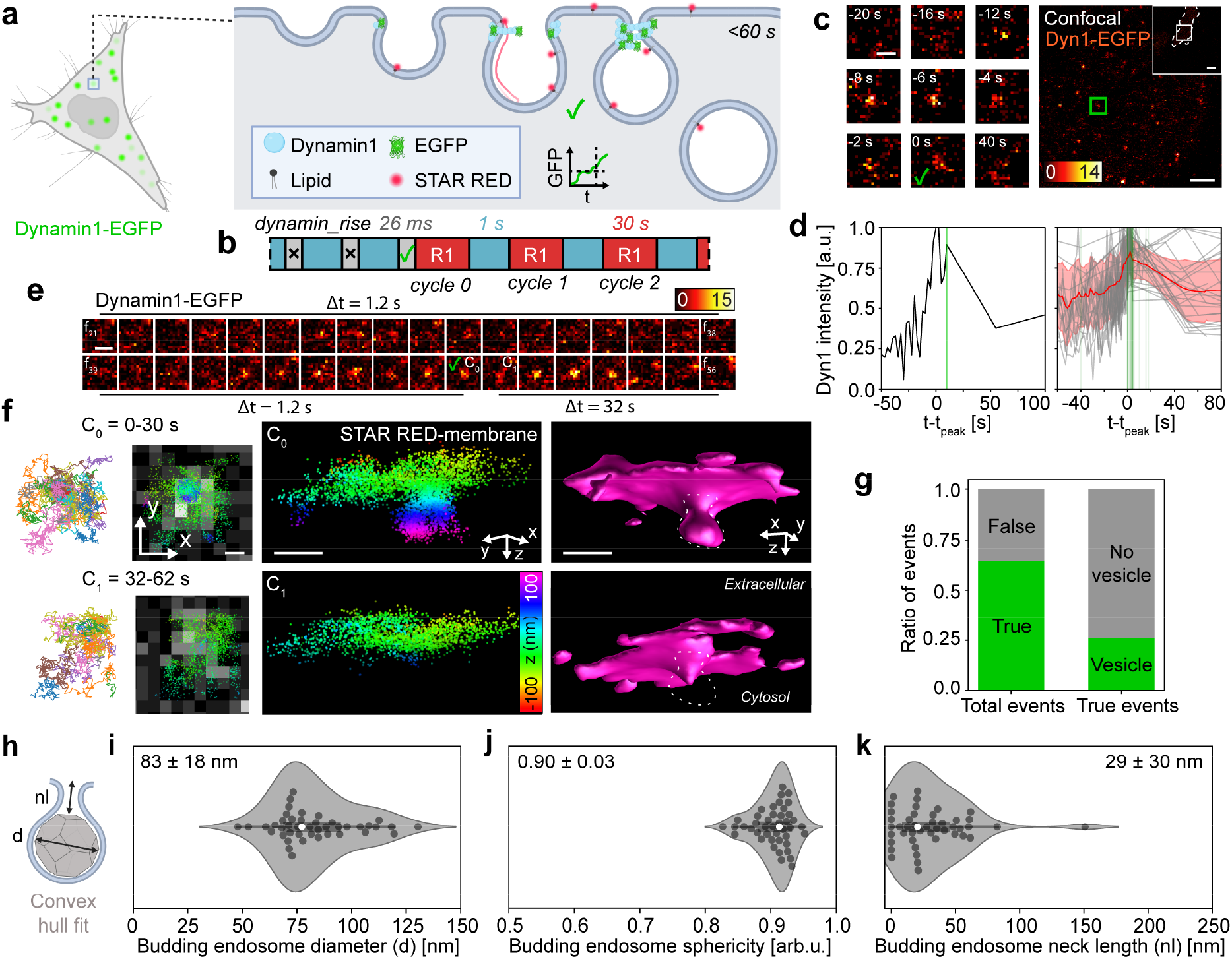
Live imaging of budding endocytic vesicles in 3D following event detection of Dynamin1 accumulation. **a**. Event progression sketch with Dynamin1 accumulation, including experimental Dynamin1-EGFP and lipid labelling with STAR RED-membrane. **b**. etMINFLUX experiment timeline, with confocal (blue), analysis (light gray), and MINFLUX (red) blocks. Indicated times refer to mean (analysis) or median (confocal, MINFLUX) times over all experimental replicas. **c**. Example of event detection, with large confocal image showing Dynamin1-EGFP (right) and zoom-ins to the event area in a time lapse collage (left). Frame triggering event is marked with a check mark. **d**. Exemplary Dynamin1 intensity event detection curve (left). All recorded event intensity curves (gray), with a mean curve (red line) with respective standard deviation (red shaded area) (right). Green lines indicate times of event detection and initiated MINFLUX tracking. **e**. Exemplary confocal zoom time lapse to the area of an event detection, showing initial 1/1.2 Hz imaging, event detection (green check mark), and 1/32 Hz imaging interleaved with MINFLUX tracking. Frames 21–56 in the confocal timelapse are shown. C_i_ indicates frame after which MINFLUX cycle i was recorded. **f**. Exemplary MINFLUX data of the event shown in **e**, showing tracks of STAR RED-membrane (first column), MINFLUX localizations overlaid on confocal zoom-in of event area (second column), 3D localization point cloud with color-coded z coordinate (third column), and surface fitting to the 3D localization point cloud (fourth column) with indicated endosome area highlighted (dashed line). Data is shown for cycle 0 (top row) and cycle 1 (bottom row) for the same event. **g**. Statistics on number of true event detections (green) as per post-experimental classification, compared to all detected events, and number of detected events with budding vesicles visible in the MINFLUX data (green), compared to all true event detections. **h**. Sketch of convex hull fitting to endosome visible in MINFLUX point cloud data, indicating how diameter (d) and neck length (nl) have been measured. **i**. Budding endosome diameter analysis. **j**. Budding endosome convex hull sphericity analysis. **k**. Budding endosome neck length analysis. **i–k**. Each datapoint represents a visible endosome before scission in a single cycle of the MINFLUX data in a true event detection. Scale bars: 10 µm (overview in **c**), 2 µm (large confocal in **c**), 500 nm (zoom-ins in **c**; confocal timelapse in **e**), 100 nm (MINFLUX-confocal overlay, 3D MINFLUX localizations, and 3D MINFLUX surface in **f**).

A typical event detection is shown in Fig. 4c. Here, a larger ROI of 15×15 µm^2^ was continuously recorded, and an event was detected at timepoint *t = 0*, where the Dynamin1 signal was continuously increasing (Fig. 4d). Following event detection, after the first MINFLUX acquisition cycle, the Dynamin1 signal had dropped, indicating the dynamin having dispersed again and an endosome likely having been released. The mean of all temporal fluorescent Dynamin1 signal curves at detected events (bold red line in Fig. 4d) indicated a rapid signal increase in the 20 s leading up to the peak Dynamin1 signal, after which the signal more slowly decayed as the Dynamin1 constriction is disassembled and the protein dispersed again. The full process finished on the scales < 60 s. Event detections were mostly done at least 5 s before the peak detected signal (green vertical lines in Fig. 4d), and likely more before the actual Dynamin1 accumulation peak due to the decreased sampling of the Dynamin1 signal after event detection and during the first MINFLUX acquisition cycle. This indicated the possibility to most commonly catch an endosome before scission in the first MINFLUX acquisition cycle.

Visualizing the data for an exemplary event, we noted a continuously increasing Dynamin1-EGFP signal appearing from a background in the confocal data, and after some time following event detection the signal decreased and disappeared, indicating a scission event (Fig. 4e). The recorded MINFLUX data corroborated the confocal data, showing a budding endosome at a late stage in the first recording cycle of 30 s (Fig. 4f, top) but a flatter membrane without an endosome head already in the next recording cycle (Fig. 4f, bottom). The presence of Dynamin1-EGFP signal for another three recording cycles may indicate that the budding endosome was either still attached but so restricted that no STAR RED-membrane probe could diffuse into it during the following recording cycles, or that the Dynamin1 took an unusual long time to diffuse away from the site. We measured many similar events, where a majority showed temporally overlapping Dynamin1-EGFP signal disappearance and 3D MINFLUX topography budding endosome disappearance (Suppl. Fig. 12).

In total, 65% of all detected events were confirmed to be real budding events in post-experimental analysis (Fig. 4g). This relatively low figure owes to the need to capture the process in an early stage, thereby omitting some frame-consuming checks that could have avoided noisy detections during the experiment. Post-experimental analysis allowed straightforward scrutinization of events thanks to the rich information and data saved for each event. Of the real events detected, only a minority of 26% showed the presence of budding endosomes in the MINFLUX data. This may be explained in different ways: (1) due to the sparse labelling necessary for MINFLUX, the fluorescent membrane probe did occasionally not enter a budding endosome present either by chance or by exclusion before scission; (2) an event was detected at a late stage of the budding and the MINFLUX acquisition was thus started too late, i.e. after endosome scission occurred; or (3) no budding vesicle was ever present. Nevertheless, the smart method allows high-throughput 3D measurements of naturally occurring endosomal events, which otherwise is difficult considering its rarity and difficulty to measure. A majority (54%) of budding endosomes visible in the MINFLUX data were present only in the first acquisition cycle (20–30 s), occasionally (15%) the same budding endosomes were present also in the second cycle (∼40–60 s) or in the third or further on cycles (15%) (∼60+ s) before seemingly undergoing scission. Rarely (15%) an identified endosome stayed for many acquisition cycles and therefore minutes following an event detection, which then indicated an arrested endosomal budding state with dynamin accumulation (Suppl. Fig. 12).

The MINFLUX topographic mapping of the captured budding endosomes allowed us to accurately measure endosomal geometry. For this, the localizations representing the endosomal surface was recognized and separated from the rest of the localization cloud using dedicated filtering steps, a convex hull was fitted to the remaining localizations, and the average position of the membrane was determined through a Gaussian fitting of the z-profile (Fig. 4h, Suppl. Note 7–8). From this, the endosomal equivalent spherical diameter (Fig. 4i), sphericity (Fig. 4j), and neck length (Fig. 4k) was extracted (Suppl. Note 7). The endosomal equivalent spherical diameter spanned 50–130 nm, with a mean size of 83 ± 18 nm, matching values derived from electron microscopy^42^. The sphericity showed almost exclusively nearly spherical endosomal heads, with a mean sphericity value of 0.90 ± 0.03. Lastly, the neck length was measured to be 29 ± 30 nm on average, with a large span of values also comprising an example of an endosome with a neck length of 151 nm. (Suppl. Fig. 13).

### EtMINFLUX 3D tracking measures membrane fluidity at Gag accumulation sites

We next tested our etMINFLUX 3D tracking procedure to monitor molecular membrane diffusion and organization at viral budding sites, taking Gag proteins as an example. Gag proteins are essential structural proteins for viruses such as retroviruses and specifically HIV. They assemble into membrane-bound aggregates from individual subunits in the early stages of virus budding at the plasma membrane of infected host cells, which is a driving factor in the bending of the membrane^44^. We used etMINFLUX with 3D MINFLUX tracking with an octahedral TCP of the fluorescent cholesterol-based membrane probe STAR RED-membrane to investigate the membrane shape and fluidity over time during the Gag accumulation process in live HeLa cells (Fig. 5a). Gag accumulation leading to virus formation and budding is a slower process than endosome budding, where the accumulation is known to take place over approximately 10 minutes^45,46^, and the full process to the time of release around 25 min^47^. As such, we applied either the single ROI follow or the multiple ROI follow mode to follow the evolution of Gag accumulation sites in both the confocal and MINFLUX channels in multiple recording cycles (Fig. 5b). We used a lower confocal frame rate around 1/20 Hz in an area of 10–25×10–25 µm^2^. MINFLUX data was acquired with recording windows of 30–60 s in around 0.7×0.7 µm^2^ ROIs. The measurements were done either at room temperature, or at 33–37 °C. Still, we present the results altogether, and the diffusion analysis is always presented as ratio values between diffusion coefficients instead of absolute values to allow comparability in the broad range of the observed absolute values of diffusion coefficients (Suppl. Fig. 14). The localization precision was determined to be 8.4 nm in these experiments (Suppl. Fig. 7). The distributions of values of the temporal track length (51 ± 84 ms), time between localizations (545 ± 873 µs, min 357 µs, median 359 µs), and localization brightness (182 ± 90 kHz) obtained from the MINFLUX metadata of a representative raw dataset are plotted in Suppl. Fig. 5.

**Figure 5.**
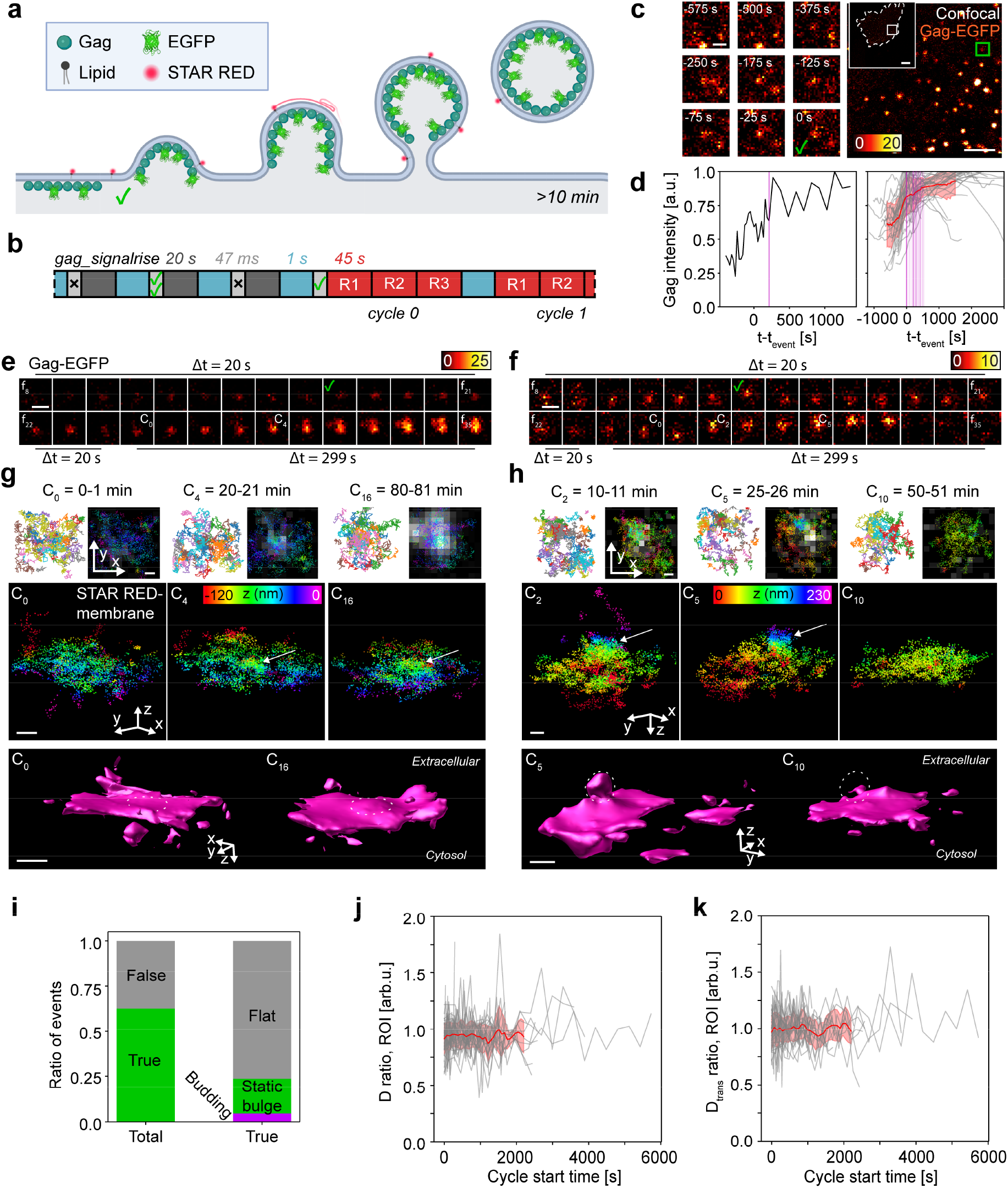
Live imaging of Gag accumulation sites and membrane fluidity in 3D following event detection of Gag accumulation. **a**. Event progression sketch with Gag accumulation, including experimental Gag-GFP and lipid labeling with STAR RED-membrane. **b**. etMINFLUX experiment timeline, with confocal (blue), analysis (light gray), and MINFLUX (red) blocks. Indicated times refer to mean (analysis) or median (confocal, MINFLUX) times over all experimental replicas. **c**. Example of event detection, with large confocal image showing Gag-EGFP (right) and zoom-ins to the event area in a time lapse collage (left). Frame triggering event is marked with a check mark. **d**. Exemplary Gag intensity event detection curve (left). All recorded event intensity curves (gray), with a mean curve (red line) with respective one standard deviation (red shaded area) (right). Magenta lines indicate start times of the first MINFLUX cycle. **e–f**. Exemplary confocal zoom time lapses to two areas of event detections, showing initial 1/20 Hz imaging, event detections (green check marks), and 1/299 Hz imaging interleaved with MINFLUX tracking. Frames 8–35 in the confocal timelapses are shown. C_i_ indicates frame after which MINFLUX cycle i was recorded. **g–h**. Exemplary MINFLUX data of the respective events shown in **e-f**, showing three recording cycles (**g**: cycle 0, 4, and 16, **h**: cycle 2, 5, and 10) of the same event. Each cycle shows tracks of STAR RED-membrane (top row, left), MINFLUX localizations overlaid on confocal zoom-in of event area (top row, right), and 3D localization point cloud with color-coded z coordinate (second row). MINFLUX data of two timepoints are shown as surface fitting to the 3D localization point cloud (third row), with areas of bulging (**g**) and budding (**h**) indicated (dashed line). **i**. Statistics on number of true event detections (green) as per post-experimental classification, compared to all detected events, and number of detected events with either a static bulging membrane (green) or a dynamically budding membrane (magenta) visible in the 3D MINFLUX tracking data, compared to all true event detections. **j**. Diffusion coefficient ratio analysis, for ratio between diffusion inside and outside the budding site, over time since MINFLUX initiation for all followed events (gray) and their mean (solid red line) and one standard deviation (shaded red area) curves. **k**. Transient diffusion coefficient ratio analysis, for ratio between diffusion inside and outside the budding site, over time since MINFLUX initiation for all followed events (gray) and their mean (solid red line) and one standard deviation (shaded red area) curves. Scale bars: 10 µm (overview in **c**), 2 µm (large confocal in **c**), 500 nm (zoom-ins in **c**; confocal timelapse in **e**,**f**), 100 nm (MINFLUX-confocal overlay, 3D MINFLUX localizations, and 3D MINFLUX surface in **g**,**h**).

Figure 5c depicts a typical event detection. A larger ROI of 10 × 10 µm^2^ was here recorded at a rate of 1/25 Hz, and an event was detected at timepoint *t = 0* after a slow and steady increase of the Gag signal over hundreds of seconds (Fig. 5d). Following the event detection and the start of MINFLUX acquisition, the confocal Gag signal kept increasing, indicating an early event detection well before the peak of Gag accumulation. This general trend in the Gag signal over time is exemplified by the mean of all temporal Gag signal curves (bold red line in Fig. 5d), where the Gag signal keeps increasing well into 30 min following event detections for most cases. In some cases, the Gag signal decreased within the 10–20 min following an event detection.

Two of the developing events are shown in Figure 5e,g and Figure 5f,h respectively. One of these events, which was recorded at the basal membrane of the cell, showed a continuously increasing Gag signal (Fig. 5e), while the other event, recorded at the apical membrane, showed accumulation of Gag signal until a point of sudden disappearance (Fig. 5f). The MINFLUX data of the first event was recorded over a total of 20 recording cycles spaced 5 min apart, and in 3D initially showed a flat membrane which then continuously bulged with an increasing size (Fig. 5g). The latter event with a suddenly disappearing Gag signal instead showed a budding and scission process, where cycles 0–2 highlighted the start to bulging (Suppl. Fig. 15a), cycle 5 showed a fully formed virus site with a narrowed neck, and cycles 7 and later, when the Gag signal had disappeared, showed a flat membrane (Fig. 5h). In total, the majority of Gag accumulation events were recorded at the bottom membrane and did not indicate any signs of bulging, budding, or Gag signal disappearance linked to virus particle scission (Suppl. Fig. 15b). A minority of events showed signs of static bulging (Suppl. Fig. 15c) and only very few showed developing budding over time (Suppl. Fig. 15d).

Our complete analysis of the event data indicated that 63% of all detected events were true events, i.e. with a continuous Gag protein accumulation, while the rest mainly consisted of fluctuations in the Gag signal that were falsely considered as an event but disappeared quickly afterwards, such as from Gag accumulation site movement (Fig. 5i). This highlights how difficult it would be to manually detect and pick out these events in real time during microscope control, and how important the automatic event-triggered control of etMINFLUX is. Of the true detected events, a clear majority of 76% showed a flat and non-changing membrane throughout the recorded cycles, while only 24% showed some bulging at the membrane. A great majority of these showed no change over recording cycles, and only 3 detected events rendered signs of budding, here defined as a membrane bulge dynamically increasing over time.

We further investigated the general membrane fluidity at the Gag accumulation sites, using the diffusion of the cholesterol-derived STAR RED-membrane a measure. We defined a diffusion coefficient ratio as the ratio between the diffusion coefficient inside and outside of the site as defined by a circle with radius 200 nm (Suppl. Note 5, Suppl. Fig. 14), calculated its value over all MINFLUX recording cycles for all true detected event sites, and aligned the resulting values to the start of each MINFLUX recording (Fig. 5j). The value of the ratio was on average around 0.9 and hardly changed over time relative to the start of the event detection, indicating a slightly lower fluidity at the Gag accumulation sites than outside of them. We also defined a transient diffusion coefficient ratio, which was calculated over all MINFLUX cycles as the ratio between the transient diffusion coefficient inside the site, as defined by a circle with radius 60 nm, and outside of a larger surrounding area, as defined by a circle with radius 240 nm (Suppl. Note 5, Suppl. Fig. 6, Suppl. Fig. 14, Fig. 5k). Compared to the firstly derived diffusion coefficient ratio, the transient diffusion coefficient ratio thus compares transient diffusion between a more centered and more distant area. Neither this ratio showed any changes over time but its mean values was closer to 1, i.e. highlighting a comparable mobility inside and outside the site. We unfortunately did not observe enough events of bulging and developing sites to investigate the fluidity at these sites as compared to Gag accumulation sites showing no change to membrane bulging.

## Discussion

We have developed a smart event-triggered approach, denoted etMINFLUX, for recording MINFLUX microscopy single-particle tracking data, which can straightforwardly also be extended to a MINFLUX imaging mode. With etMINFLUX we extend the current developments in smart microscopy with the state-of-the-art super-resolution microscopy performance of MINFLUX. We applied etMINFLUX to three different biological systems and cellular events, which proves its versatility. For more static event sites such as caveolae we highlighted that etMINFLUX can increase the throughput and gather data from hundreds of sites with minimal human interaction; for a rapid event process such as that of budding endosomes we depict that etMINFLUX consistently enables to capture cellular events on time scales down to tens of seconds even in 3D that would be unfeasible with manual control; and for more slowly developing event sites such as Gag accumulation we show that etMINFLUX allows to detect the sites at an early stage and follow the development in 3D at multiple sites simultaneously, once again significantly increasing the throughput of the data collection. To our knowledge, this is the first time budding endosomes and viruses have been imaged in 3D and with such high spatial resolution in living cells under physiological conditions, proving the power of the smart etMINFLUX method and topographic data extracted from 3D MINFLUX tracking. Overall, this proves etMINFLUX as a method for expanding the capabilities of MINFLUX to systems where it was previously inapplicable. The implementation of the method that controls a commercial abberior MINFLUX system, and is further released open source, will importantly enable a wide range of users to take advantage of the method.

## Methods

### MINFLUX microscope

All experiments were performed on a commercial abberior MINFLUX microscope (abberior Instruments GmbH, Göttingen, Germany). Details of the microscope and components are previously fully described^5^. Importantly, the microscope is built around an inverted microscope body (IX83, Olympus, Hachioji, Japan). The lasers used in the presented experiments were: 488 nm (confocal), 642 nm (MINFLUX). In the presented experiments, one stabilization system was used in a feedback loop that stabilizes the sample in all three spatial dimensions. The stabilization system uses a 980 nm IR laser and an xyz sample piezo (Piezoconcept). The objective used was a 100x 1.45 NA UPLXAPO oil objective (Olympus). Fluorescence detection was performed in two spectral channels for the MINFLUX measurements using two APDs, 650–685 nm and 685–750 nm, from which the signal was summed, and one spectral channel for the confocal measurements using one APD, 500–550 nm. The microscope was controlled using the Imspector control software (v16.3.15635, abberior Instruments). The microscope was aligned before each measurement session using a gold beads and fluorescent beads sample (abberior Instruments).

A stage-top incubator with an objective heater (Microscope Heaters, Brighton, UK) was used in order to set a higher temperature in the experiments presented in Fig. 5e–h and Suppl. Fig. 15a, c–d, reaching sample temperatures of 34–35 °C. All other presented experiments were performed at room temperature. For the analysis results on the Gag data experiments presented in Fig. 5, a mix between room temperature and incubated experiments were performed. No difference between the two groups was observed. When using an objective heater, the microscope was aligned after the incubator and alignment sample reached a stable high temperature.

### etMINFLUX control widget

The etMINFLUX control widget (further explained in Suppl. Note 1) is a self-standing control widget written in Python, based on the design of the control widget previously developed for event-triggered STED imaging^22^ in the ImSwitch environment^48^. The etMINFLUX control widget is entirely self-standing and open source. It is available on GitHub (https://github.com/jonatanalvelid/etMINFLUX), and the version used at the submission of this manuscript, v1.0.0, is available on Zenodo (https://doi.org/10.5281/zenodo.16967461). The control widget interfaces with the commercial Imspector control software using the *specpy*^49^ Python package, as well as Python packages *mouse*^50^ and *pynput*^51^ for simulated mouse and keyboard control respectively. The control widget has been tested with Imspector version v16.3.15635 and specpy v1.2.3 on a commercial abberior MINFLUX setup, running on Windows. Other than standard and aforementioned Python packages, the widget has been implemented using the *pyqtgraph, PyQt5*^52^, *matplotlib*^53^, *tifffile*^54^, *scipy*^55,56^, and *numpy*^57^ packages.

### MINFLUX tracking sequences

Custom-adjusted sequences were used for MINFLUX tracking, based on the default fast triangle 2D and octahedron 3D tracking sequences provided by abberior Instruments. The tracking sequences were optimized for each experimental condition, and kept consistent throughout all experiments with the same sample type. The most important sequence parameters are listed in the table below. For all three sequences, the stickiness was set to 4 and the damping was set to 1. Any other parameters were not changed from their default values. The three sequences are provided as supplementary material. Resulting metadata value distributions (temporal track length, *dt*, and *efa*) are representatively shown for the three different experiment types in Suppl. Fig. 5.

| Iteration | Pattern | TCP L (nm) | Photon limit | Dwell time (μs) | Pattern repeats | CFR limit | Background threshold (kHz) | Laser power factor |
| --- | --- | --- | --- | --- | --- | --- | --- | --- |
| **seqTrk2D-cav-SR: Caveolae (Caveolin1), 2D tracking, SR-SM and SR-DPPE** |  |  |  |  |  |  |  |  |
| 0 (Preloc.) | Hexagon | - | 20 | 500 | 1 | - | 15 | 1 |
| 1 | Hexagon | 302 | 10 | 100 | 1 | 0.5 | 30 | 1.5 |
| 2 | Hexagon | 151 | 6 | 100 | 1 | - | 30 | 2 |
| 3 | Triangle | 100 | 15 | 50 | 1 | - | 74 | 3 |
| **seqTrk3D-dyn-SRmemb: Endocytosis (Dynamin1), 3D tracking, SR-membrane** |  |  |  |  |  |  |  |  |
| 0 (Preloc.) | Hexagon | 288 | 40 | 500 | 1 | - | 30 | 1 |
| 1 | z-line | 288/1440 | 300 | 2000 | 1 | - | 30 | 1 |
| 2 | Octahedron | 288 | 40 | 200 | 1 | - | 40 | 1 |
| 3 | Octahedron | 151 | 30 | 200 | 1 | 0.9 | 55 | 2 |
| 4 | Octahedron | 100 | 20 | 200 | 1 | - | 71 | 3 |
| **seqTrk3D-gag-SRmemb: Virus budding sites (Gag), 3D tracking, SR-membrane** |  |  |  |  |  |  |  |  |
| 0 (Preloc.) | Hexagon | 288 | 40 | 500 | 1 | - | 30 | 1 |
| 1 | z-line | 288/1440 | 300 | 2000 | 1 | - | 30 | 1 |
| 2 | Octahedron | 288 | 40 | 200 | 1 | - | 40 | 1 |
| 3 | Octahedron | 151 | 30 | 200 | 1 | 0.9 | 50 | 2 |
| 4 | Octahedron | 100 | 20 | 200 | 1 | - | 60 | 3 |

### Confocal and MINFLUX acquisition parameters

Full confocal and MINFLUX acquisition parameters for all data presented in this work are available in Suppl. Table 2. The ranges of confocal and MINFLUX acquisition parameters were the following: 488 nm confocal excitation 0.6–3.0 µW; 642 nm MINFLUX excitation 39–41 µW; MINFLUX ROI sizes 0.6–2×0.6–2 µm^2^; MINFLUX acquisition times 20–90 s; confocal fast axis scanning speeds: 3.5–50 µm/ms; confocal pixel sizes 70–100 nm.

### Cell culture

HeLa (ATCC CCL-2) cells were cultured in DMEM (9007.1, Carl Roth GmbH + Co. KG, Karlsruhe, Germany) supplemented with 10% (vol/vol) fetal bovine serum (10082147, Gibco, Thermo Fisher Scientific, Waltham, USA) and 1% penicillin-streptomycin (15140122, Gibco, Thermo Fisher Scientific), and maintained at 37 °C and 5% CO2 in a humidified incubator. Cells were seeded on 8-well glass bottom µ-slides (80827 or 80807-90, ibidi GmbH, Gräfelfing, Germany) 24 h (Gag) or 48 h (Dynamin1) before imaging.

PtK2 (ATCC CCL-56) cells were cultured in DMEM (9007.1, Carl Roth) supplemented with 15% (vol/vol) fetal bovine serum (10082147, Gibco, Thermo Fisher Scientific) and 1% penicillin-streptomycin (15140122, Gibco, Thermo Fisher Scientific), and maintained at 37 °C and 5% CO2 in a humidified incubator. Cells were seeded on 8-well glass bottom µ-slides (80827 or 80807-90, ibidi) 48 h before imaging.

### Transfections

For transfections, cells were transfected 1 day (18–24 h) after seeding using lipid-nanoparticle-based transfection reagent BioT (B01, Bioland Scientific LLC, Paramount, USA) or Lipofectamine 3000 (L3000015, Invitrogen, Thermo Fisher Scientific), according to the instructions of respective manufacturer. When transfecting with Gag plasmids, both plasmids were used with a ratio of Gag-wt 1:10 Gag-EGFP to ensure a properly functioning Gag lattice formation^58,59^. Transfection times and plasmid amounts were optimized experimentally, qualitatively judging transfection efficiency and cell health with brightfield, widefield, and confocal imaging. Optimized transfection times were 15 h (Dynamin1), 6 h (Gag), and 15 h (Caveolin1). Optimized plasmid amounts were 81 ng/well (Dynamin1), 100 ng/well (Gag), and 125 ng/well (Caveolin1).

### Dye-labeled lipid analogues

Lipid labelling was performed using one of the following dye-labeled lipid analogues: STAR RED-membrane (cholesterol functional group, STRED-0206, abberior GmbH, Göttingen, Germany), STAR RED-C12 sphingosyl PE (SM, STRED-0201, abberior), or STAR RED-1,2-dipalmitoyl-sn-glycero-3-phosphoethanolamine (DPPE, STRED-0200, abberior).

### Caveolae experiments sample labeling

The PtK2 cells were labelled using STAR RED-SM at a concentration of 0.1–0.9 nM or STAR RED-DPPE at a concentration of 0.8–2.5 nM in Leibovitz L-15 medium (21083027, Thermo Fisher Scientific), with the concentration optimized at each experiment day to ensure that the MINFLUX tracking was following single dyes. The cells were immediately after labeling put on the microscope and imaged for a maximum time of 1 h, while ensuring that that MINFLUX tracking was following single dyes during the full experiment time.

### Dynamin1 and Gag accumulation sites sample labeling

The HeLa cells were labelled using STAR RED-membrane at a concentration of 150–300 pM for Dynamin1 experiments or 100– 300 pM for Gag experiments, with the dye in L-15, and with the concentration optimized at each experiment day to ensure that the MINFLUX tracking was following single dyes. The cells were incubated with the dye for 5 min at 37 °C, and then washed 2× with L-15. Fresh L-15 was then added to the wells as imaging medium.

### Plasmids

Cav1-GFP was a gift from Ari Helenius (Addgene plasmid #14433; http://n2t.net/addgene:14433; RRID:Addgene_14433)^60^. pEGFP-N1 Dynamin1 wt was a gift from Justin Taraska (Addgene plasmid #120313; http://n2t.net/addgene:120313; RRID:Addgene_120313)^61^. Gag-EGFP and Gag wt were gifts from Jennifer Lippincott-Schwartz.

### Gold beads labeling

For use with the XYZ sample stabilization system of the MINFLUX microscope, 150 nm gold colloid particles (EM.GC150/4, BBI Solutions, Crumlin, United Kingdom) were added to each sample. At the beginning of the sample preparation protocol, ∼30 min before experiment start and ∼15 min before lipid dye labeling, a solution of 120 µl stock solution in 400–450 µl L15 imaging medium was added to the cells. The gold bead solution was left to incubate for 15 min at 37°C, followed by washing 6–8× with 250 µl L15 imaging medium.

### Confocal-MINFLUX scan shift compensation

Coordinate system scan shifts between confocal and MINFLUX data were observed and compensated for according to necessity in post-processing of the acquired data. This was applied for all Caveolin1 event data. The procedure for measuring and applying shift compensation is further explained in Suppl. Note 2.

### Diffusion and transient diffusion analysis

Diffusion analysis was performed using square displacement analysis and subsequent fitting of the population of displacements and time steps with a function representing a free Brownian motion in two dimensions, i.e. on the membrane. A 2D Brownian diffusion model in a limited time step span, using a maximum time lag of 500 µs or 5 ms for the 2D and 3D MINFLUX tracking diffusion analysis respectively, was always assumed as we analyzed the short-range diffusion of lipid analogues in the membrane to investigate the immediate diffusion at sites never larger than a few hundred nanometers. From each fit, the diffusion coefficient and dynamic localization precision could be extracted. Diffusion analysis was performed on whole sections of tracks, while transient diffusion analysis was performed for each localization using a sliding window centered on the localization. For further details of the diffusion analysis, see Suppl. Note 5.

### Caveolae sites analysis results filtering

Thanks to the relatively large amount of data recorded, the final analysis results considered for the caveolae sites were filtered according to the following considerations in order to ensure robust results. For diffusion coefficient analysis, only those track sections where the number of square distances was above 15 were considered, and only those sites where the number of fitted diffusion coefficients, i.e. number of track sections, inside and outside were both above 5 were considered. Furthermore, the mean diffusion coefficient of the track sections inside and outside were both required to be above 0. For the transient diffusion coefficients, only the sites where the total number of tracks was above 20 and where the number of localizations inside was above 50 were considered. For the inclusion ratio T, only regions of interests recorded with the same ROI size, 1 x 1 µm2, and where the total number of tracks was above 20 were considered.

### Data visualization

*3D surface fitting*. For 3D surface visualization of MINFLUX tracking data we use the *ch-shrinkwrap*^62^ localization cloud surface fitting plugin to PYME^24^. To generate a shrinkwrap surface, an isosurface is first fitted to the localization cloud. This is performed with following parameter value ranges: N points min, 12–20; threshold density, 0.00001–0.0001; inner surface culling, remeshing, and smooth curvature were used. Following this, the shrinkwrap algorithm is performed on the created isosurface, with the following parameter values or value ranges: curvature weight, 7; remesh frequency, 5; neck first iteration, 5; neck threshold low, -0.00003–-0.001; neck threshold high, 0.01; punch frequency, 0; finishing iters, 0; Kc, 0.9; minimum edge length, 5.0; truncate at, 1000; smooth curvature was used. *3D point cloud z-threshold*. In the following 3D point cloud visualizations, as well as overlay between confocal and a 2D projection of the localization point cloud, a z-threshold has been used to only extract the data from the membrane on which the event of interest is occurring: Fig. 5g, Fig. 5h, and Suppl. Fig. 15a. On all of these occasions, both cellular membranes have been captured and are present in the raw 3D MINFLUX tracking, and the threshold has been introduced manually for each case in order to allow for accurate analysis and clearer visualization.

### Localization precision determination

The localization precision was chosen to be extracted as the dynamic localization precision from fitting of the square displacement analysis, rather than subsequent step distance analysis due to the diffusional nature of the tracked molecules. With the linear fitting model used (Suppl. Note 5), the diffusion coefficient can be extracted as the slope while the dynamic localization precision can be extracted as the intersection with the vertical axis. All tracks were fitted individually, the dynamic localization precision was extracted from each track where the number of square distances from the track was above 15, and the individual datapoints in Suppl. Fig. 7 represents the mean of all tracks in a region of interest around a detected event site.

### Statistics

Statistical tests are either one-sample t-tests for the null hypothesis that the mean is equal to 1 (Fig. 3f,g); or two-sample t-tests for the null hypothesis that the two independent (Fig. 3h,i) or related (Suppl. Fig. 8) populations have identical means. Statistical significance is presented in the following way: n.s., p > 0.05; ^*^, p < 0.05; ^**^, p < 0.01; ^***^, p < 0.001.

### Experimental sizes

Total number of experiment days: 5/5/3/3 (Caveolin1: SM-Cav1/DPPE-Cav1/SM-random/DPPE-random), 4 (Dynamin1), 12 (Gag). Total number of samples: 13/14/5/5 (Caveolin1: SM-Cav1/DPPE-Cav1/SM-random/DPPE-random), 10 (Dynamin1), 35 (Gag). Total number of cells: 17/30/7/7 (Caveolin1: SM-Cav1/DPPE-Cav1/SM-random/DPPE-random), 33 (Dynamin1), 42 (Gag). Total number of detected events: 101/141/31/29 (Caveolin1: SM-Cav1/DPPE-Cav1/SM-random/DPPE-random), 155 (Dynamin1), 106 (Gag).

## Data availability

The data supporting the method implementation and supporting the findings in this study, including images, log files, and metadata, are available at Zenodo, reference number 15608840^63^.

## Code availability

The source code of the developed control widget in this study is open source and available on GitHub: https://github.com/jonatanalvelid/etMINFLUX and in a released version as the version used for the data collection in this manuscript, v1.0.0, at Zenodo, reference number 16967461^64^. The code of the analysis supporting the findings in this work, in the forms of Jupyter notebooks, Python scripts, and Python class modules, together with example data, is available on GitHub: https://github.com/jonatanalvelid/etMINFLUX-analysis-public and Zenodo, reference number 16967494^65^.

## Author contributions

C.E. supervised the research. C.E. and J.A. funded the research. J.A. and C.E. conceived the project idea. J.A. and C.E. designed the research. J.A. planned and developed the control widget and the real-time analysis pipelines; planned, prepared, and performed experiments; and performed data analysis. A.K. planned, prepared, and performed experiments; and provided experimental and data analysis guidance. J.A. drafted the manuscript with input from all the authors. All authors revised and accepted the final version of the manuscript.

## Acknowledgements

J.A. thanks the Swedish Research Council (2022-06139_VR) for supporting this work. The authors thank the Deutsche Forschungsgemeinschaft (DFG, German Research Foundation; Instrument funding MINFLUX Jena INST 275_405_1) for supporting this work. The authors thank Annett Urbanek for help with sample preparation and the Microverse Imaging Center (and Aurélie Jost / Patrick Then) for providing general microscope facility support for data acquisition (and data analysis). The authors acknowledge further financial support by the Deutsche Forschungsgemeinschaft (Germany’s Excellence Strategy – EXC 2051 – Project-ID 390713860; project number 316213987 – SFB 1278; GRK M-M-M: GRK 2723/1 – 2023 – ID 44711651; GRK PhInt: GRK 3014/1; instrument funding ID 460889961 multi-photon laser scanning device), the State of Thuringia (TMWWDG), the Leibniz Association (Leibniz Collaborative Excellence Programme, project AMPel – project number K548/2023), and the Free State of Thuringia (TAB; Advanced Flu-Spec / 2020 FGZ: FGI 0031; Multi-XUV / 2023 FGR 0054). Further, this work is supported by the BMFTR (Federal Ministry of Research, Technology and Space), funding program LIVE2QMIC (FGZ: 13N15956) as well as Photonics Research Germany (FKZ: 13N15713 / 13N15717) and is integrated into the Leibniz Center for Photonics in Infection Research (LPI). The LPI initiated by Leibniz-IPHT, Leibniz-HKI, UKJ and FSU Jena is part of the BMFTR national roadmap for research infrastructures. A part of the project on which these results are based was funded by the Free State of Thuringia under the number 2018 IZN 0002 (Thimedop) and co-financed by funds from the European Union within the framework of the European Regional Development Fund (EFRE). Figures 3a (https://BioRender.com/4rus5v5), 4a (https://BioRender.com/ah6nktp), 4h (https://BioRender.com/rzieic9) and 5a (https://BioRender.com/f6vehwf) were created with BioRender; Alvelid, J. (2025).

