## Supplementary Information for "Event-triggered MINFLUX microscopy: smart microscopy to catch and follow rare events"

#### List of content

1. **Supplementary Notes**
2. **Supplementary Figures**
3. **Supplementary Tables**
4. **Supplementary References**

#### List of Supplementary Notes


1. EtMINLUX widget
2. Scanning-dependent confocal-MINFLUX coordinate shifts
3. Real-time analysis pipelines
4. MINFLUX track data filtering steps in caveolae analysis
5. Diffusion analysis on 2D and 3D MINFLUX tracking data
6. Simulated Brownian motion and MINFLUX tracking
7. Geometrical analysis of endosomal vesicles
8. Z-artefact filtering of 3D MINFLUX tracking data


#### List of Supplementary Figures


1. EtMINFLUX widget GUI
2. Further experimental modalities enabled by event-triggered MINFLUX
3. Scanning-dependent confocal coordinate shifts
4. Analysis pipeline performance
5. Exemplary dataset metadata value distributions
6. Caveolin site data analysis sketch
7. MINFLUX localization precision
8. Diffusion analysis on simulated data
9. Diffusion analysis results from Caveolin1 accumulation sites
10. etMINFLUX caveolae and random sites with SM-STAR RED – further examples
11. etMINFLUX caveolae and random sites with DPPE-STAR RED – further examples
12. etMINFLUX endocytosis sites – further examples
13. etMINFLUX endocytosis site with long endosome neck
14. Diffusion coefficient and transient diffusion coefficient analysis results from Gag accumulation sites
15. etMINFLUX virus budding sites – further examples


#### List of Supplementary Tables


1. etMINFLUX analysis pipeline parameter value ranges
2. Confocal and MINFLUX acquisition parameters in the various experiments

### Supplementary Notes

#### Supplementary Note 1. EtMINFLUX control widget

The etMINFLUX control widget is a software module written in Python using a view-controller design pattern, and is based on the generic event-triggered microscope control widget previously developed as part of event-triggered STED imaging[1]. The control widget was developed to be self-standing and interacts with the Abberior MINFLUX microscope control software Imspector. The widget is entirely open source and available on GitHub.

The control widget interacts with Imspector using three means: the *specpy* package[2], which is a Python-based API for Imspector released by Abberior Instruments; the *mouse* package[3], which allows user-like simulation of mouse movements; and *pynput*[4], which allows user-like simulation of keyboard interaction. The reason for the need of the latter two is missing commands in the *specpy* API for MINFLUX-specific controls, due to a lack of a package version after MINFLUX microscope release. An integration of new methods in the API would allow *mouse* and *pynput* to be removed and allow for a more user-friendly widget implementation.

The control widget is built up of individual subwidgets, allowing the user to control what and how information is displayed during etMINFLUX experiments: the main widget, a calibration widget, an event detection overview widget, and an event view widget. The main widget has controls for the real-time analysis pipeline, the experiment mode in use, how the real-time analysis is run, the MINFLUX acquisition, the binary mask recording, and various data saving modalities. The real-time analysis pipeline control starts with choosing which pipeline to load, after which all the defined parameters for the pipeline shows up in the GUI with editable fields. This allows the user to tweak the parameter values to the sample and experiment at hand. Before running an experiment, the user can choose to record a binary mask using the positive and negative thresholds and smoothing options. The binary mask will be sent to each analysis pipeline call and can be applied in order to limit the area that is considered for the analysis.

After setting the analysis pipeline parameters, the recording mode of the etMINFLUX method can be set using two separate controls. Firstly, the user can change between experiment, visualization test, and validation test modes. The former runs the experiments as normal, while the latter two are two different modes that can be used while not triggering MINFLUX acquisitions. Here, the confocal imaging is continued as normal and the analysis pipeline is run, but as soon as an event is detected, the pipeline either continues to run the confocal as normal while in visualization mode, or the experimental event data is saved, including a set of confocal images that are recorded after the event is detected, which allows the user to further validate the event detection in post-acquisition analysis. In the visualization mode, the detected events are marked on top of the latest confocal image in the event detection overview widget, allowing the user to easily see what is being detected as events and allows a fast feedback for optimizing and tweaking the analysis pipeline parameters. In this widget, a list of all the events with their pixel positions and region of interest (ROI) IDs is further shown. For normal experiment runs, the second set of recording mode controls allows the user to control the ROI following modes. Here, the user can activate ROI following, i.e. interleaved MINFLUX and confocal measurements of an event site after event detection, and chose the type of ROI following mode: single-site, multi-site, single-site with redetection, and multi-site with multi-detection. Two further parameters can be set: the MINFLUX acquisition time per ROI per cycle, as well as a redetect distance threshold that will be used in the redetection mode that will keep running the analysis pipeline in the interleaved confocal images after event detection and will update the event site position for each MINFLUX cycle. The latter recording mode is particularly useful for an event site that can move over time.

The analysis control part of the widget further allows control over how the analysis pipeline will be run. The base case is to run it after each recorded confocal frame, however here it can optionally be set to run after a set number of lines, the analysis period, if faster event detection is required. Furthermore, a pause between confocal frames can be set, if slower event appearance is expected; a number of initial frames to ignore can be set; and an option to show the event detection overview widget while running the experiment mode can be set. In ROI following modes the latter defaulted to be on and allows direct interaction with the list of event ROIs where they are continuously sorted according to the next ROI in line to be recorded, in the case of multiple ROI following, and individual ROIs can be deleted to allow full focus on the events of interest.

Lastly, before running an experiment, the MINFLUX acquisition parameter can be set from the etMINFLUX main widget. As MINFLUX acquisition parameters otherwise requires to enter the MINFLUX mode of Imspector, these cannot be set preemptively by the user directly in Imspector. Instead, the MINFLUX sequence, the MINFLUX excitation laser and excitation power, potential activation power, as well as ROI size and acquisition time can all be set in the main etMINFLUX widget. The corresponding values in Imspector will automatically be set by the widget before a MINFLUX acquisition takes place, which also allows the user to tweak these between cycles in the ROI following mode. Boolean controls if to record ROIs in random positions of the binary mask (i.e. random instead of event sites); if to trigger subsequent MINFLUX acquisition of all detected events; and if to use a pre-set ROI size and MINFLUX acquisition time or not are also provided – if not the ROI size needs to be determined by the analysis pipeline, and the MINFLUX acquisition time will be indefinite.

When initiating an etMINFLUX experiment, the experiment can be run in endless mode, which will repeat the process after finishing a MINFLUX recording, allowing the widget to control the automatic experiments indefinitely. When an event recording has finished, the MINFLUX, confocal, and metadata will be automatically saved. An option is provided if the user does not want to automatically save the .msr file with the MINFLUX data, and furthermore an option to automatically delete MINFLUX datasets after an event recording can also be found. When an event is detected, the etMINFLUX will automatically draw a ROI in the right place and initiate the MINFLUX recording.

The calibration subwidget hosts options about the screen configuration, which needs to be calibrated in order for the automatic mouse movements to function properly. Here the user needs to detect the top left and bottom right corners of the confocal recording window in Imspector (which needs to be unobstructed during etMINFLUX recordings), after which the widget will automatically read the confocal image size in pixels and micrometers from

Imspector, which finally allows a conversion between image position and screen position to be able to draw the MINFLUX ROIs at the right place. There is also a way to calibrate the screen positions of the repeat measurement button and the top MINFLUX datasets in the dataset viewer widget in Imspector, which will be used to click the button and select the top MINFLUX dataset for deletion (if that option is selected) respectively. These three types of functionality are not available in the *specpy* Python API for Imspector and thus needs to be handled by simulated moues movements and clicks. In the calibration subwidget there are further parameters for various timing options that are needed for proper functioning due to the simulated mouse movement interactions necessary – if drawing the ROI too fast or if clicking the necessary buttons too fast after each other the control is unreliable. For optimal temporal performance, these timings need tweaking between individual microscope control computers as they depend on the computer specifications.

The last subwidget is the event view, that is shown when a MINFLUX acquisition is initiated at an event site. In this subwidget, the user can view a zoomed-in view of the event site over the whole timelapse stack of confocal images recorded. Additionally, a graph is shown plotting the confocal intensity summed in a small area around the event site over time. A dropdown menu allows the user to switch between all detected event sites, and the stacks and graphs are continuously updated during a ROI following experiment, allowing the user to inspect and decide if to keep or delete ROIs before further cycles are recorded.

#### Supplementary Note 2. Scanning-dependent confocal-MINFLUX coordinate shifts

Despite being scanned by the same scanning apparatus and sharing a global coordinate system, there are residual shifts between the confocal and MINFLUX coordinate systems that needs to be considered for optimal overlapping of confocal and MINFLUX data. This is not applicable only to etMINFLUX data, but any confocal and MINFLUX data recorded on the Abberior MINFLUX microscope. The shifts we have characterized depend on the scanning speed of the confocal image, where we can approximate the galvanometric mirrors to be stationary during MINFLUX acquisition. The shift for a specific point in the image depends also on the distance from the start of a scanning line. The shifts occur both in the X and Y direction, with significantly different types of shift visible. While we have not understood the underlying sources of the full range of shifts we observe, we have fitted compensation curves to test measurements allowing us to correct for it in post-processing of data. Optimally, a slower confocal scanning is used, in which case no significant shift is observed and a compensation can be left out. Based on our testing, we have deemed a confocal linear scanning speed of 7 µm/ms (pixel size 70 nm, pixel dwell time 10 µs) safe to avoid causing shifts in need of compensation in our final data. In the Caveolin1 data, we have compensated for the shifts present, while in the Dynamin1 and Gag data we have used a slower confocal scanning speed.

The scanning-dependent coordinate shifts were measured by using confocal imaging with two different scanning speeds – very slow, to mimic a pseudo-stationary state as used during MINFLUX acquisition, and the scanning speed of choice in the experiments. The very slow speed was set to 0.7 nm/µs. With these settings, two confocal images were taken subsequently of a test sample containing fluorescent beads. Each bead position in the image was automatically detected in the two images, the beads found in both images were connected, and the position of them were fitted to subpixel precision using 2D gaussian fitting. The center of each peak was extracted, and the shift in X, fast scanning axis, and Y, slow scanning axis, for each bead in the two images was calculated. This shift can be assumed to be identical to the shift that would occur in a MINFLUX acquisition following a fast confocal acquisition, which was confirmed by compensating the shift in confocal-MINFLUX measurements of fluorescent beads (data not shown) with the shifts extracted from confocal-confocal measurements. The shift population can be plotted in a 2D scatter plot, where each datapoint is an individual bead, the horizontal axis shows the distance from the left border of the image for that bead, and the vertical axis shows the shift in either X or Y. Doing this, the spatially dependent shifts become apparent (Suppl. Fig. 3). Recording multiple instances of pairs of images, across different days, confirmed the consistent spatially-dependent shapes of the shifts. In the figure, data from multiple image pairs has been overlaid and fitted together.

The X-shift of a non-bidirectional scan with a scanning speed of 35 nm/µs (Suppl. Fig. 3a) takes the form of an initial second-degree polynomial followed by a sigmoidal function, while the Y-shift (Suppl. Fig. 3b) takes the form of an oscillating sinusoidal function with an overall downwards slope. In a bidirectional scan with the same scanning speed, the X-shift (Suppl. Fig. 3c) takes the shape of a linearly increasing function, while the Y-shift (Suppl. Fig. 3d) takes the shape of a linearly decreasing function. Finally, the shifts from a bidirectional scan with a different scanning speed (50 nm/µs) are shown (Suppl. Fig. 3e–f), confirming that the shape stays the same but the slope changes for different scanning speeds. This is also valid for non-bidirectional scans (data not shown).

The large range of shift values measured for a specific position from the leftmost part of the image can be explained by random scan-to-scan shifts that cannot be compensated for. Overall, the average scan shifts measured, which is what the compensation is based on, can be up to 150 nm large, depending on the scanning speed. The residual random and uncorrectable shifts can be determined from the plots to not be larger than ~50 nm, and hence is the residual error after correction. Knowing this, a slower scanning speed to avoid the shifts is preferable to compensating for the shifts in post-processing of the data.

### Supplementary Note 3. Real-time analysis pipelines

The etMINFLUX widget uses real-time analysis pipelines that shares a common structure, which allows them to be flexibly loaded in the widget, and allows users to develop new analysis pipelines that can be directly used for experiments. The pipelines are developed as Python functions with a specific initial call syntax, including the stack of previously recorded images, a potential binary mask, a fed-back Python object of any type, and an end call syntax that contains all the pipeline parameters that the user can tweak in real-time in the etMINFLUX main widget after loading the pipeline.

Three different real-time analysis pipelines were used during the process of this work, each one optimized at the task at hand in the three different application cases presented. While the pipelines share common features, it is important to note that each pipeline should be optimized to the specific sample at hand. The below-presented pipelines have each been optimized, methodologically and for parameter values, on training data recorded as confocal timelapses without etMINFLUX activated. These timelapses have been manually annotated as a ground truth for the events one would wish to detect, to which the outcome of the real-time analysis pipelines has been compared and further optimized. Each of the three pipelines are presented in detail below.

#### Real-time peak detection analysis pipeline

The *peak_detection* real-time analysis pipeline was used in testing and characterization, in order to detect fluorescent beads, as well as in detection Caveolin1-EGFP accumulation sites, i.e. caveolae. The pipeline is supplemented to the etMINFLUX control widget as a separate Python module containing only the pipeline function. It uses only the last confocal image as input, and outputs the coordinates of any detected peaks, according to the pipeline parameters provided by the user. The pipeline has default values for all input parameters, and the full parameter value ranges that were used for the data recording in this work are presented in Supplementary Table 1. The pipeline uses the *numpy*[5], *scipy*[6,7], *scikit-image*[8], and *opencv*[9] packages. The pipeline is available on GitHub:
https://github.com/jonatanalvelid/etMINFLUX/blob/main/analysis_pipelines/peak_detection_bright.py.

The pipeline starts with a smoothing with a Gaussian filter, using the user-provider provided smoothing radius. The pipeline can take a binary mask as input, which if provided is used to mask the image after smoothing. Following this, the image is dilated, with a rectangular kernel of a user-provided size. The dilated image is compared to the original smoothed image, and any pixels with an equal value in the two images, and a peak value above a user-provided threshold, are considered peaks. The peaks are sorted according to pixel intensity, and any peaks inside a border limit of a user-provided size are removed. Furthermore, peaks that are too close to each other are removed, as they will be considered to be in a too crowded area. The distance between peaks that are considered too close is user provided. In the end, only the first N peaks in the list are returned, allowing the user to chose how many peaks at most they want.

Following the peak detection, if the user has requested local ROI sizes to be calculated, a further part of the analysis pipeline will take place. This part looks at the peak coordinates, and for each peak it gets a mask of the immediate surrounding of all pixels above a certain user-provided percentage of the peak value. From the mask, the x and y sizes of the smallest bounding box of the largest area in the mask will be calculated. These will be returned as the ROI size.

Multiple version of the pipeline exists, where the returned peak coordinates are sorted or filtered in a different way depending on the need: (1) brightest peaks first, (2) dimmest peaks first, (3) only the peak of a random peak index, or (4) only the peak of a user-provided peak index.

#### Dynamin rising signal analysis pipeline

The *dynamin_signalrise* real-time analysis pipeline was used to detect dynamin1-EGFP rising over time on the seconds timescale, indicating an accumulation of dynamin1 and an endocytosis scission site. The pipeline is supplemented to the etMINFLUX control widget as a separate Python module containing only the pipeline function. It uses the last confocal image as well as a stack of the previously recorded frames as input, and outputs the final coordinates of any spot that has been considered as having a rising signal according to the conditions and pipeline

parameters explained below. The pipeline has default values for all input parameters, and the full parameter value ranges that were used for the data recording in this work are presented in Supplementary Table 1. The pipeline uses the *numpy*[5], *scipy*[6,7], *opencv*[9], *pandas*[10], and *trackpy*[11] packages. The pipeline is available on GitHub:
https://github.com/jonatanalvelid/etMINFLUX/blob/main/analysis_pipelines/dyn_signalrise.py.

The pipeline starts with a smoothing with a Gaussian filter of a fixed radius of 1.5 pixels. The image is then fed through two separate gaussian filters, with a 0.05- and 3-pixel radius respectively, and a difference of Gaussians image is calculated between the two. This step is in order to make the signal peaks clearer and more separated from spread-out background and noise in the image. Negative pixel values are at this point clipped and set to 0. Following this, a third step of Gaussian filtering is performed, once again with a 1.5-pixel radius.

After this, a peak detection is performed in the following steps: firstly, a rectangular kernel is determined using a user-inputted parameter as the size; secondly, the image is dilated with this kernel; thirdly, a mask containing the pixels that are equal in the dilated image and the image before dilation is calculated. Two thresholds, one high and one low, provided by the user are used to filter the mask based on the pixel intensities in the image prior to dilation of the peaks that are represented in the mask. This removes any detected peaks not inside the intensity range of interest. The coordinates of the peaks are put in a list and are sorted in a descending order according to their pixel intensities. The list is also cut after a number of peaks corresponding to a user-inputted parameter, for control over how many peak tracks to follow. The intensity summed in a 5-pixel-wide rectangle around each detected peak is extracted, and the timepoint, coordinates, and intensities are all put into a *pandas* dataframe. This dataframe is concatenated with the information from all previously recorded frames and runs of the pipeline, which is returned and inputted to each pipeline run.

After this, the conditional checking of tracks is initiated, to find the spots that increases in intensity. First, the dataframe is provided to the *link* method in *trackpy*, that performs a single-particle track linking algorithm and finds the most likely connected tracks from the peaks detected in each frame, using user-inputted parameters for the search range in time and space. After this, five checks are performed on each track, and an event is detected only if it passes all conditional checks. Check 1 checks if a track appeared, i.e. had the first timepoint of the track in the dataframe a certain number of frames ago, where the number of frames is provided by the user. Check 2 checks that the track stayed for most frames after that, and only allows it to be gone for one frame. Check 3 checks that the intensity of the track increase over time from appearance to the current frame. It does this by getting the mean of the intensity before appearance at the spot of appearance, the intensity track around appearance, and the intensity track after appearance, where each window is a few frames long, controlled by user-inputted parameters. The means are then turned into two intensity ratios, comparing the around appearance with before appearance, and the after with around appearance. Both of these ratios have to be inside a threshold range, controlled by two inputted parameter values, to be considered valid – above a threshold to actually be a true increasing signal and below a threshold to avoid detecting rapidly appearing signal that likely indicates a lot of movement or noise. Check 4 checks that the track has not moved too much since appearance, and does so by taking the mean Euclidean distance the spot has moved between frames since appearance, and compares this mean to a user-provided threshold. This again ensures a decrease in false detections as the dynamin should accumulate at a relatively fixed point on the membrane. Finally, check 5 checks that the last position of the track is not at the border of the image, with the border size decided by a user-inputted parameter, as this otherwise hinders the MINFLUX ROI from being set correctly. If all conditional checks are passed, the event coordinates are extracted and the pipeline is immediately returned, as we are not interested in any potential additional sites as the endocytosis happens too fast to be able to catch multiple scission events simultaneously.

#### Gag rising signal analysis pipeline

The *gag_signalrise* real-time analysis pipeline was used to detect Gag-EGFP rising over time on the minutes timescale, indicating an accumulation of Gag and a potential virus budding site. The pipeline is supplemented to the etMINFLUX control widget as a separate Python module containing only the pipeline function. It uses the last confocal image as well as a stack of the previously recorded frames as input, and outputs the final coordinates of any spot that has been considered as having a rising signal according to the conditions and pipeline parameters explained below. The pipeline has default values for all input parameters, and the full parameter value ranges that were used for the data recording in this work are presented in Supplementary Table 1. The pipeline uses the *numpy*[5], *scipy*[6,7], *opencv*[9], *pandas*[10], and *trackpy*[11] packages. The pipeline is available on GitHub:
https://github.com/jonatanalvelid/etMINFLUX/blob/main/analysis_pipelines/gag_signalrise.py.

The image preprocessing starts with a Gaussian smoothing step with a 1.5-pixel radius kernel, followed by a peak detection similar to the peak detection described for the dynamin-detecting pipeline: dilation with a rectangular kernel, comparison between dilated and before-dilation images, and extraction of pixels with an equal intensity. The peak coordinates are filtered with a high and low threshold, sorted according to intensity, and everything beyond a certain number of peaks is removed. After this, a *pandas* dataframe with the time, coordinates, and intensity in 5-pixel-wide rectangle is defined, and added onto any previous data that is refed into the pipeline with every run.

After this, the conditional checking of tracks is initiated, to find the spots that increases in intensity. First, the dataframe is provided to the *link* method in *trackpy*, that performs a single-particle track linking algorithm and finds the most likely connected tracks from the peaks detected in each frame, using user-inputted parameters for the search range in time and space. After this, seven checks are performed on each track, and an event is detected only if it passes all conditional checks. Check 1 checks if a track appeared inside 1–3 × *frames_appear* ago, where *frames_appear* is a user-inputted parameter – here a range of frames are considered as it increases the chance that an event does not go undetected due to noisy intensity traces. Check 2 checks that the peak stays detected for at least 70% of the frames following the appearance, allowing some flickering in peak detection during the early stages of the site where the intensity is close to the threshold. Check 3 checks if the distance to all other peaks during appearance is above a user-provided threshold, which ensures that no events are detected due to the tracks of two or more peaks crosses each other and the become visible as multiple peaks again. Check 4 checks that the intensity of the peak increases over time. This check is here performed differently depending on the length of the frames available and number of frames since the track appeared. If there are not enough frames available, similar intensity ratio checks between before, during, and after appearance as described for the *dynamin* detection above are performed, while if more information is available, a curve fitting takes place to be more precise and robust to noise. A linear function is fitted to the track intensity over time, and the slope of the fit is extracted. If the slope is above a user-provided threshold (usually between 4–10%/frame), or in the previous case if the intensity ratios are passed, check 5 is performed. This check checks that the final intensity is above a lower threshold and below an upper threshold, to avoid noisy, low-intensity detections and likely other high-intensity false detections. Check 6 looks at the mean moving distance between frames for all frames where the peak is detected, and compares this to an upper threshold. Finally, check 7 checks that the last position of the track is not at the border of the image, with the border size decided by a user-inputted parameter, as this otherwise hinders the MINFLUX ROI from being set correctly. If all conditional checks are passed, the event coordinates are extracted and the pipeline is returned.

#### Supplementary Note 4. MINFLUX track data filtering steps in caveolae analysis

Single-molecule lipid tracks from MINFLUX data were filtered before lipid diffusion analysis took place. Tracks were filtered on a track basis, where for each track three quality checks were performed. The first check was to see that the temporal track length was longer than 30 ms, in order to ensure enough statistics for better diffusion coefficient estimates in the downstream analysis. The other two checks were made to ensure that the track was not just depicting a fluorophore seemingly stuck in the same place. Depending on the probe used, we observed more or less stuck tracks, and while the source of these stuck tracks is unknown it is beyond the scope of this study to further understand them. We focused on the tracks that showed movement away from such a stuck position at some point during the track. In order to sort out these tracks, we checked that the mean distance moved between positions with a 50 localizations interval along the full track was above 13 nm, and that the mean of the standard deviations of the positions from a 40-localizations-long sliding window was above 20 nm. These thresholds were set by judging the filtering applied to test datasets, tweaking the threshold values until a balance between filtering faulty tracks and keeping diffusing and long enough tracks was found. An example of the filtering can be found in Suppl. Fig. 6, where blue indicates short tracks, gray indicates a failed mean movement distance threshold, red indicates a failed mean standard deviation of positions threshold, and green indicates that all checks were passed.

Additionally, the confocal data of the caveolae sites was used in order to ensure that the caveola at which the diffusion analysis was going to take place did not move significantly during the MINFLUX recording wherever data was available to support this analysis. In order to do this, the confocal frames before the MINFLUX acquisition, as well as after the MINFLUX acquisition in terms of the confocal image before the subsequent MINFLUX acquisition if performed in the same FOV, were extracted. Small ROIs around the detected caveolae were extracted and a 2D Gaussian function was fitted to the intensity signal. The center of the fitted Gaussian was compared before and after the MINFLUX acquisition, and if the distance between the two centers was larger than 70 nm, i.e. roughly one pixel in the confocal images, the peak was considered to have moved and no diffusion analysis was performed. In the datasets were the confocal data necessary was not available, higher intensity cav1 peaks were selected which were considered from confocal timelapses to be more stable. The fitted caveola position in the confocal frame before the MINFLUX acquisition was used as the center position of the caveola in subsequent spatial analysis.

### Supplementary Note 5. Diffusion analysis on 2D and 3D MINFLUX tracking data

In MINFLUX tracking data, the time steps ($dt$) between localizations are generally not equal due to various factors but mainly three; the target coordinate pattern can repeat due to not reaching the photon limit and will thus multiply the time step with an integer factor, there is an additional time due to calculations and preparations for the next localization, and there is temporal jitter. Together, this means that time steps are uneven and thus complicates the use of standard mean squared displacement (MSD) analysis as used in single-particle tracking with a camera, where the time steps are defined as the frame interval. Several approaches can be taken to ensure an accurate diffusion analysis from the MINFLUX data, and the one we employ here is a population-based square displacement analysis.

To perform this analysis, we start by gathering all possible time steps and their corresponding particle displacement distances ($dd$) between localizations up to a time step limit from the track or population of tracks we want to extract a diffusion coefficient for. Whereas an MSD analysis at this point would take the mean displacement for all displacements of a certain time step, and then fit a function to the means, our square displacement analysis instead fits a function directly to the 2D population of values on the form $(dt, dd)$. The function fitted is identical to that which would be used in MSD analysis. From the fit, a diffusion coefficient can be extracted, if the number of displacements on which the fit is performed exceeds a certain threshold to ensure a reliable value.

The fitting function used in this work is the following function previously introduced by Michalet and Berglund[12] and further refined by Balzarotti et al.[13] for MINFLUX analysis:

$$dd = 2dDdt + 2d\sigma^2 - 4dR_{blur}Ddt_{median},$$

where $dd$ is the displacement, $dt$ is the time step, $d$ is the number of dimensions of the diffusion model, $D$ is the diffusion coefficient, sigma is the dynamic localization precision, $R_{blur}$ is the motion blue coefficient, and $dt_{median}$ is the median value of the time step population. In the diffusion analysis performed here, $R_{blur}$ was set to 1/6.2 [13] and $d$ was set to 2, as the diffusion model assumed was always two-dimensional on a membrane even when the tracking performed was in 3D. $dt_{median}$ was set to 85.7e-6 and 350e-6 in the 2D and 3D tracking analysis, respectively.

In our analysis, we extract both diffusion coefficients and transient diffusion coefficients. The difference between these is only the population on which we perform the analysis. For diffusion coefficients, we perform the analysis on the displacement population of a full track or the sections of the track that are inside respectively outside the defined radius of the event site. For transient diffusion coefficients, we perform the analysis on the displacement population of a sliding window around the localization to which we assign the fitted diffusion coefficient.

A main limitation of the diffusion coefficient analysis applied on cases where we want to calculate the diffusion coefficient in a small site area is the bias that is introduced due to the limited area. As we require a certain number of displacements for a fit to be reliable, for a small enough radius the only track sections that will meet that requirement are those that linger for a longer time at the site, and not those that pass it faster. Thus, the diffusion coefficients inside a site will be biased towards lower diffusion coefficients. Analysis performed on simulated diffusion data (Suppl. Note 6), with a diffusion coefficient and localization uncertainty modelled after that observed in our MINFLUX tracking data, indicated that a 200 nm radius was necessary to avoid such a bias, and hence this radius was used for the analysis (Suppl. Fig. 8). The transient diffusion coefficient analysis does not suffer from such a bias, and hence a smaller radius can be used.

### Supplementary Note 6. Simulated Brownian motion and MINFLUX tracking

The simulated MINFLUX tracking data was simulated as a Brownian motion process. For each set of parameters, 10 ROIs each with 100 tracks were simulated. The track length (in number of localizations) was determined individually for each track by taking a random value from an exponential distribution with scale parameter 250, matching the distribution of the real MINFLUX 2D tracking data. For each new localization in the iterative track simulation, a random time step $dt$ is first pulled from a weighted discrete distribution matching that of real experimental data, to mimic the uneven time steps of MINFLUX data. Then, two values were added to the previous localization for each dimension: (1) a step with a random value pulled from a normal distribution with a standard deviation of $\sqrt{2Ddt}$, where $D$ is the simulated diffusion coefficient and $dt$ is the randomized time step, to simulate the molecule movement; and (2) a random value pulled from a normal distribution with a standard deviation equal to the extracted dynamic localization precision from the real MINFLUX tracking data, i.e. 9 nm, to mimic the localization process. To mimic the sites of Caveolin1 accumulation with a lower diffusion coefficient, the diffusion coefficient $D$ is dependent on the molecule position. At each new localization step, the distance to the simulated site center is checked, and depending on if it is above or below the simulated site radius, here set to 50 nm, $D_{in}$ or $D_{out}$ is used.

To mimic the MINFLUX acquisition process and potential loss of the molecule during tracking due to the molecule moving too far from the targeted coordinate pattern center: if the molecule movement step is larger than 75 nm the molecule is considered lost and the track is terminated.

### Supplementary Note 7. Geometrical analysis of endosomal vesicles

The endosomal vesicles before scission in the Dynamin1 accumulation sites data were geometrically analyzed using a pipeline centered around a convex hull fit. All parameters of the following filtering and analysis were optimized for each endosome, to ensure an accurate extraction of geometrical values. To start with, assuming a relatively flat membrane, which was the case in a vast majority of recorded data, the average membrane position was found using

Gaussian fitting of the summed z-profile. Depending on if one or both membranes were visible in the tracking data a single or a double Gaussian function was used. The peak position was considered the membrane position, and the σ of the Gaussian was considered the membrane width. Following this, all localizations of the membrane were filtered out, and only localizations at a distance of generally 1–2σ above the membrane were kept. After this, a DBSCAN clustering algorithm[14] was used to cluster the data into clusters, with each cluster ideally representing an endosome. In most cases, only one cluster was found, while in occasional cases where multiple endosomes or other noisy tracks were present a label corresponding to the endosome in question was manually chosen. A minimum number of samples of 100 and an eps of 0.04 was generally chosen for the clustering algorithm.

With the localizations of the endosome cluster, an outlier removal was performed prior to the convex hull fitting, to get a more accurate shape of the endosome that otherwise would always be overestimated. The outlier removal used the *IsolationForest* function from the *scikit-learn* package[15], using 10 estimators and contamination at 0.25. Following this, a convex hull was fitted, using the *ConvexHull* function of the *scipy* package[6,7]. The surface area $A$ and volume $V$ were extracted from the fit, and the sphericity $S$ was calculated as:

$$S = \frac{\pi^{1/3}(6V)^{2/3}}{A}.$$

The equivalent spherical diameter D was calculated as:

$$D = 2\left(\frac{3V}{4\pi}\right)^{1/3}.$$

Lastly, the neck length was calculated as the difference between membrane position in z and $D/2$ subtracted from the z-centroid of the convex hull.

For endosomal vesicles present in multiple MINFLUX acquisition cycles, the cycle with the clearest endosome visible in the data was chosen to be analyzed.

### Supplementary Note 8. Z-artefact filtering of 3D MINFLUX tracking data

A common artefact in 3D MINFLUX tracking data are z-artefacts, where the track artificially continuously seems to move far in z in the same direction localization after localization, despite this being unreasonable due to the sample constrictions (such as these tails entering into the cover glass), while having longer times between localizations, due to pattern repeats. They occur more in data with a higher fluorophore concentration, and are therefore likely connected to background signals affecting the MINFLUX localization process, further corroborated by the fact that multiple pattern repeats are needed to meet the photon limit criteria. In order to allow visualization and accurate analysis of the data, such artefacts are filtered away in any 3D MINFLUX data as a first step in the post-processing and analysis of the data.

The filtering is based on the longer time between localizations (dt) observed during the artefacts. It uses a sliding window approach, and calculates the average dt in a window of 70 localizations for each localization. For the sequences used in the experiments of this work, a dt threshold was set to 550 μs, and all localizations with a sliding window threshold above that was flagged as artefacts. The sliding window size and dt threshold need to be optimized depending on the sequence and excitation laser power used.

### Supplementary Figures

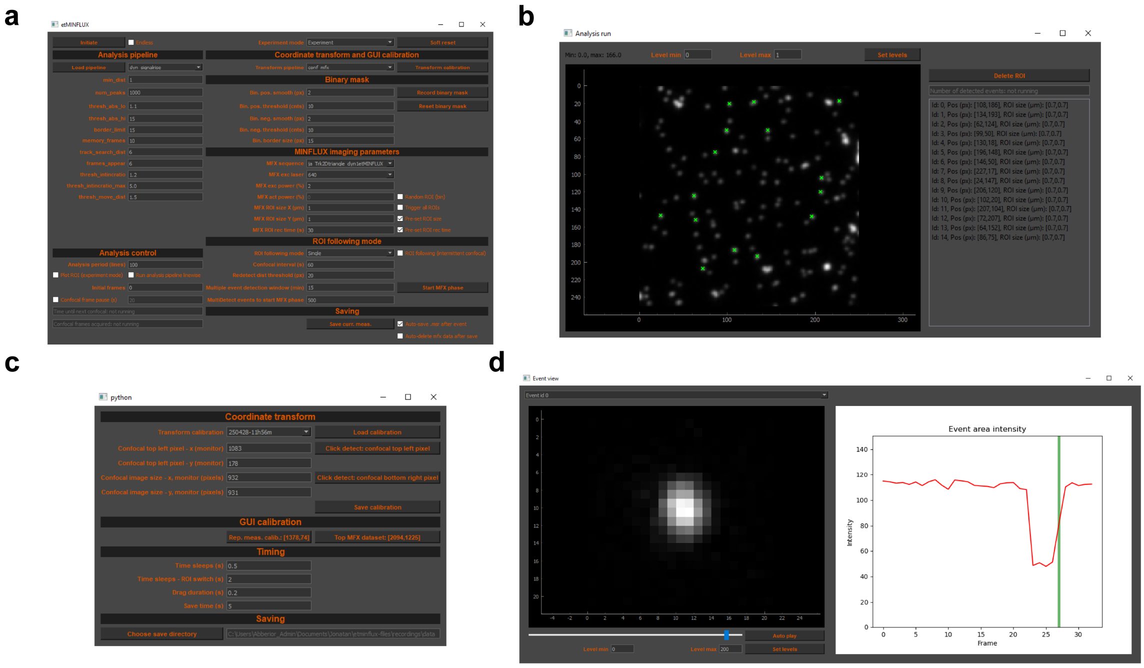


**Suppl. Figure 1. EtMINFLUX widget GUI. a.** GUI of main widget window, with analysis pipeline settings (top left), analysis control (bottom left), coordinate transform, GUI calibration, and binary mask settings (top right), MINFLUX acquisition settings (middle right), and ROI following mode and saving settings (bottom right). **b.** Analysis pipeline live-view and event detection sites window, with processed analysis image and overlaid detected events (left) and interactable list of detected events (right). **c.** Calibration settings window, with calibration settings for coordinate transform (top), Imspector GUI screen calibration settings (middle), and timing settings (bottom). **d.** Event view window, with a time lapse viewer of a zoom-in of the chosen detected event (left) and an intensity graph over the full time lapse of the immediate surrounding of the event site (right).

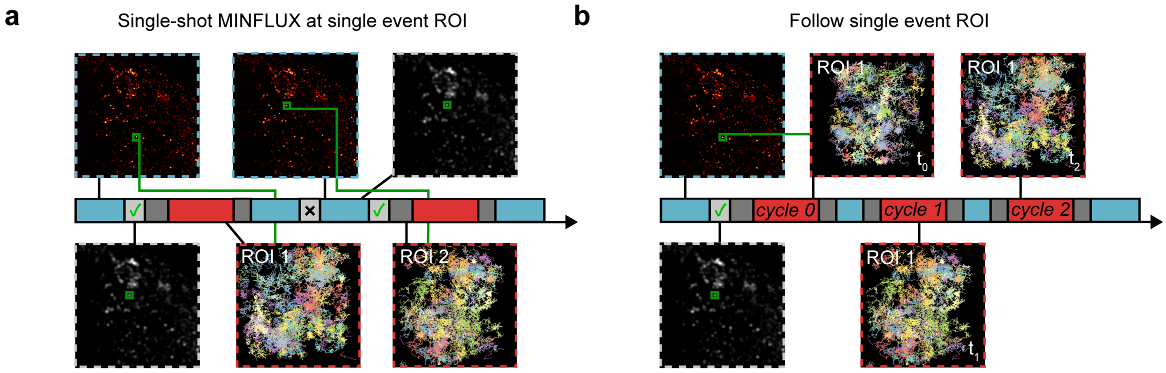


**Suppl. Figure 2. Further experimental modalities enabled by event-triggered MINFLUX. a.** Single event ROI recording after a single event. **b.** Single event ROI following, with interleaved confocal and MINFLUX recordings. Timescales show confocal acquisition (blue), MINFLUX acquisition (red), real-time analysis (light gray) and overhead (dark gray).

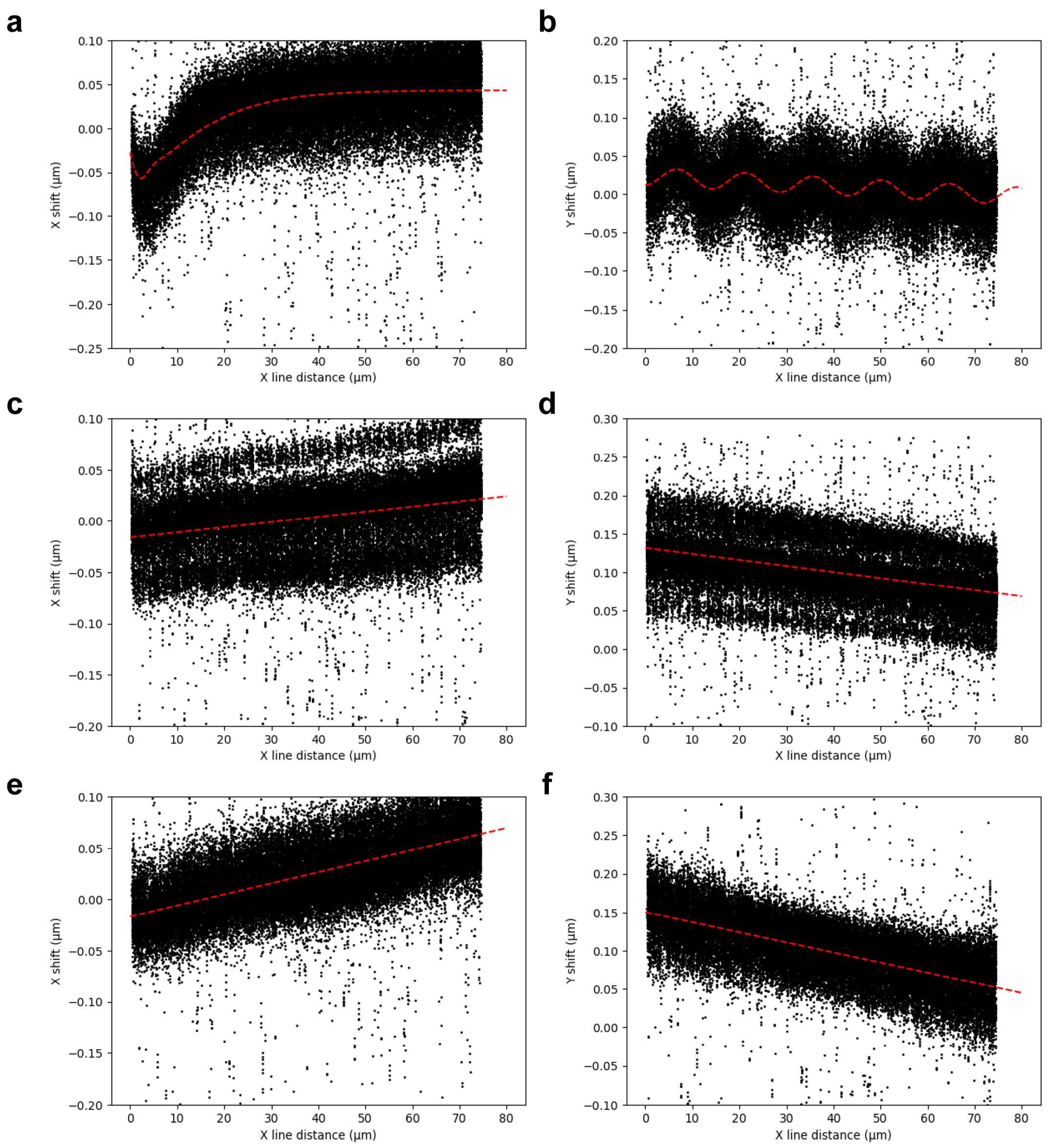


**Suppl. Figure 3. Scanning-dependent confocal coordinate shifts.** Acquisition parameter dependent coordinate shifts between fast and slow confocal image acquisitions. Each datapoint represents a bead and the measured X- or Y-axis shift (y) at the bead distance from the start of the fast-axis line (x) for a fluorescent bead in an image pair. The data from multiple image pairs have been overlapped. **a.** X-shift for unidirectional scanning with a scanning speed of 35 nm/µs. **b.** Y-shift for unidirectional scanning with a scanning speed of 35 nm/µs. **c.** X-shift for bidirectional scanning with a scanning speed of 35 nm/µs. **d.** Y-shift for bidirectional scanning with a scanning speed of 35 nm/µs. **e.** X-shift for bidirectional scanning with a scanning speed of 50 nm/µs. **f.** Y-shift for bidirectional scanning with a scanning speed of 50 nm/µs.

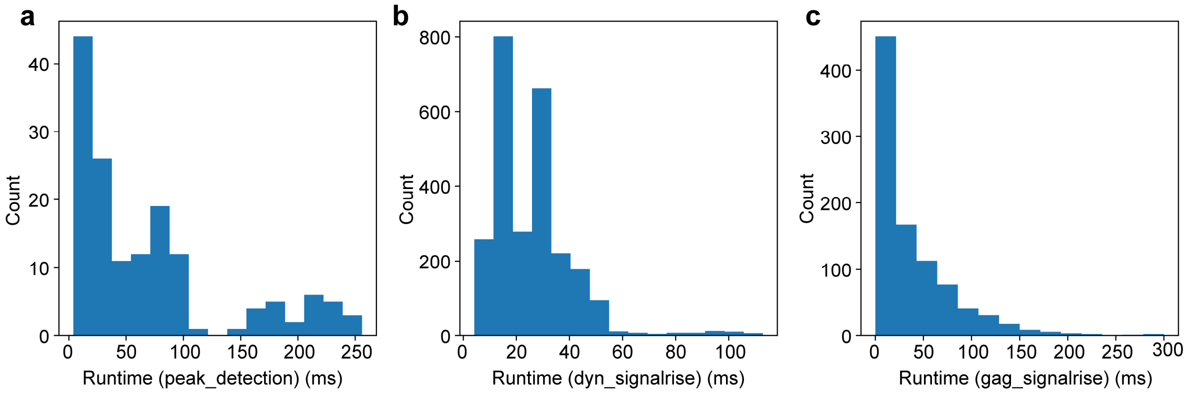


**Suppl. Figure 4. Analysis pipeline performance.** Mean (bar height) run time for analysis pipeline applied in the experiments, with one standard deviation (error bar) shown. **a.** Runtimes for *peak_detection* analysis pipeline used in caveolae site experiments. **b.** Runtimes for *dyn_signalrise* analysis pipeline used in Synamin endocytosis experiments. **c.** Runtimes for *gag_signalrise* analysis pipeline used in Gag accumulation experiments.

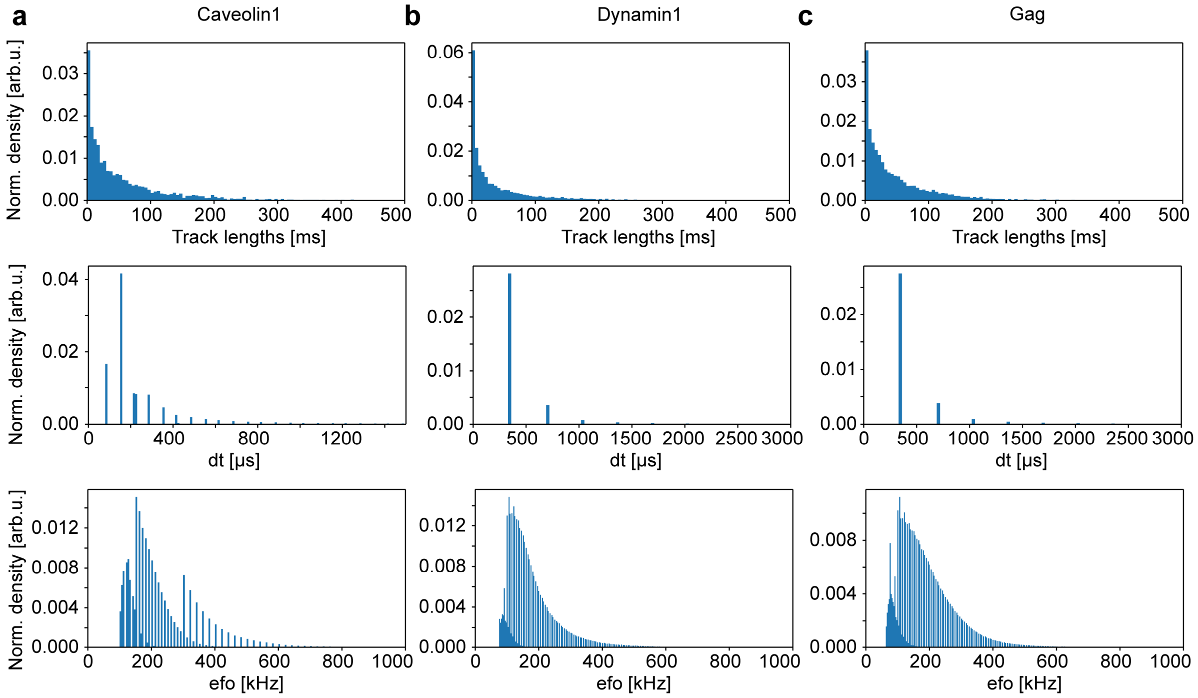


**Suppl. Figure 5. Exemplary dataset metadata value distributions.** Representative metadata value distributions for one dataset from each experiment type, showing track length, time between localizations, and efo distributions, for Caveolin dataset in **a**, Dynamin dataset in **b**, and Gag dataset in **c**.

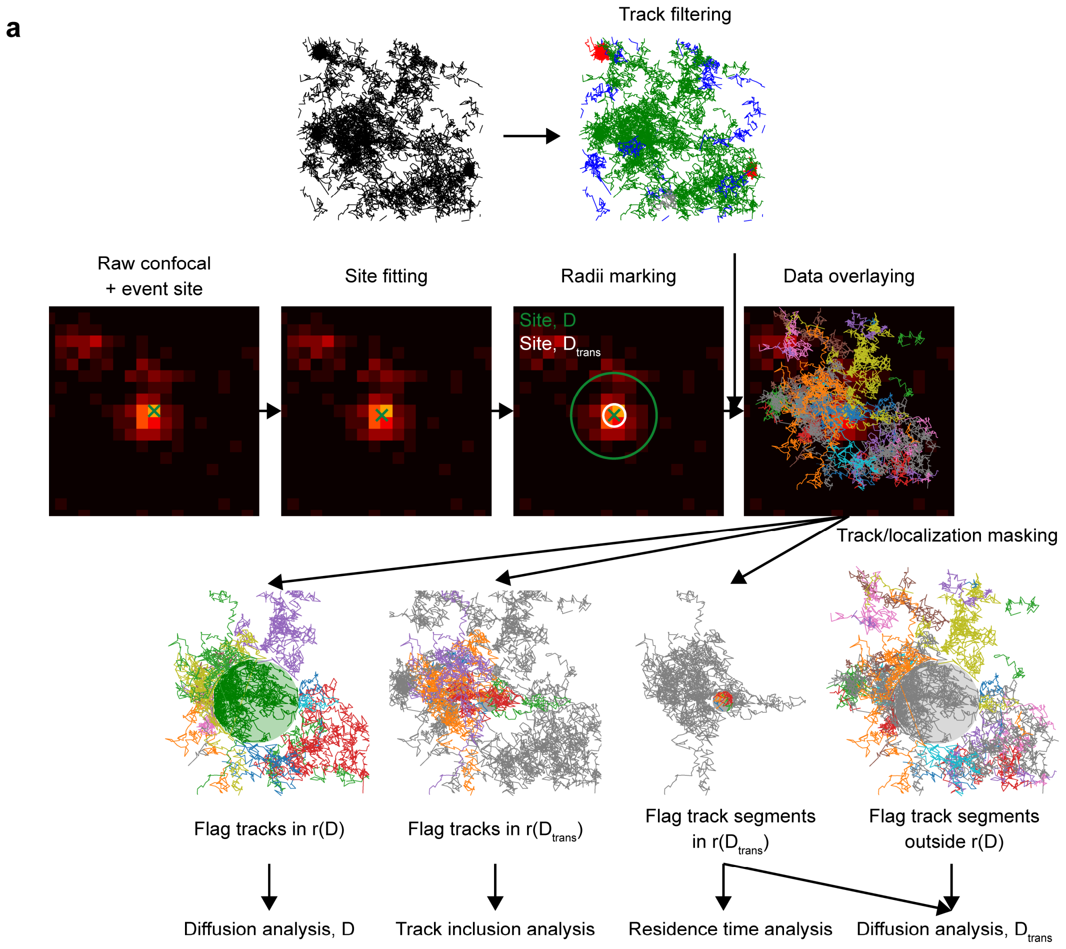


**Suppl. Figure 6. Caveolin site data analysis sketch. a.** Post-acquisition data analysis pipeline of Caveolin1 and random site MINFLUX tracking, showing steps of track filtering, site fitting, site radii marking, data overlaying, track and localization flagging, and parameter extraction for the analysis present in the work.

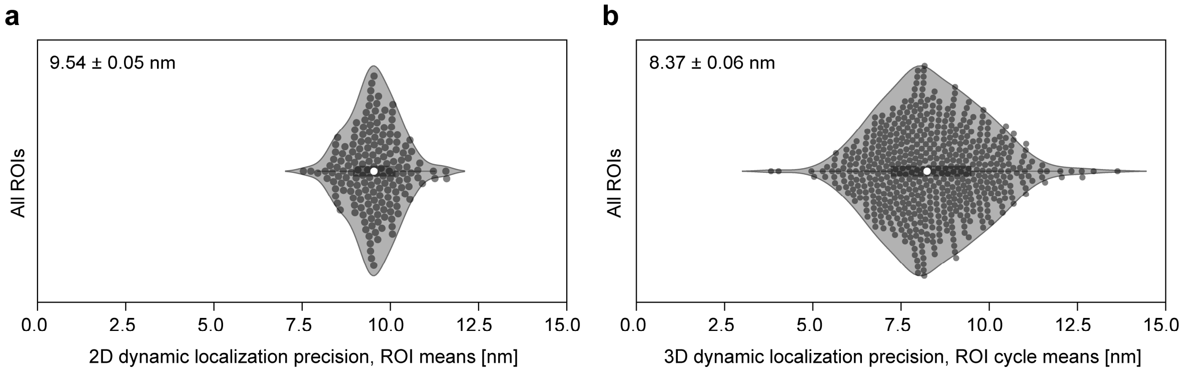


**Suppl. Figure 7. MINFLUX localization precision.** Dynamic localization precisions, extracted as the y-axis intercept from SD model fitting, for the various MINFLUX acquisition settings for the experiments on Caveolin1 accumulation sites and Gag accumulation sites. Each datapoint represents the mean of all SD fits in a ROI or ROI cycle. **a.** 2D dynamic localization precision for 2D MINFLUX tracking of SM-STAR RED or DPPE-STAR RED in Caveolin1-triggered or random site experiments (all datasets combined). **b.** 3D dynamic localization precision for 3D MINFLUX tracking of STAR RED-membrane in Gag-triggered site experiments.

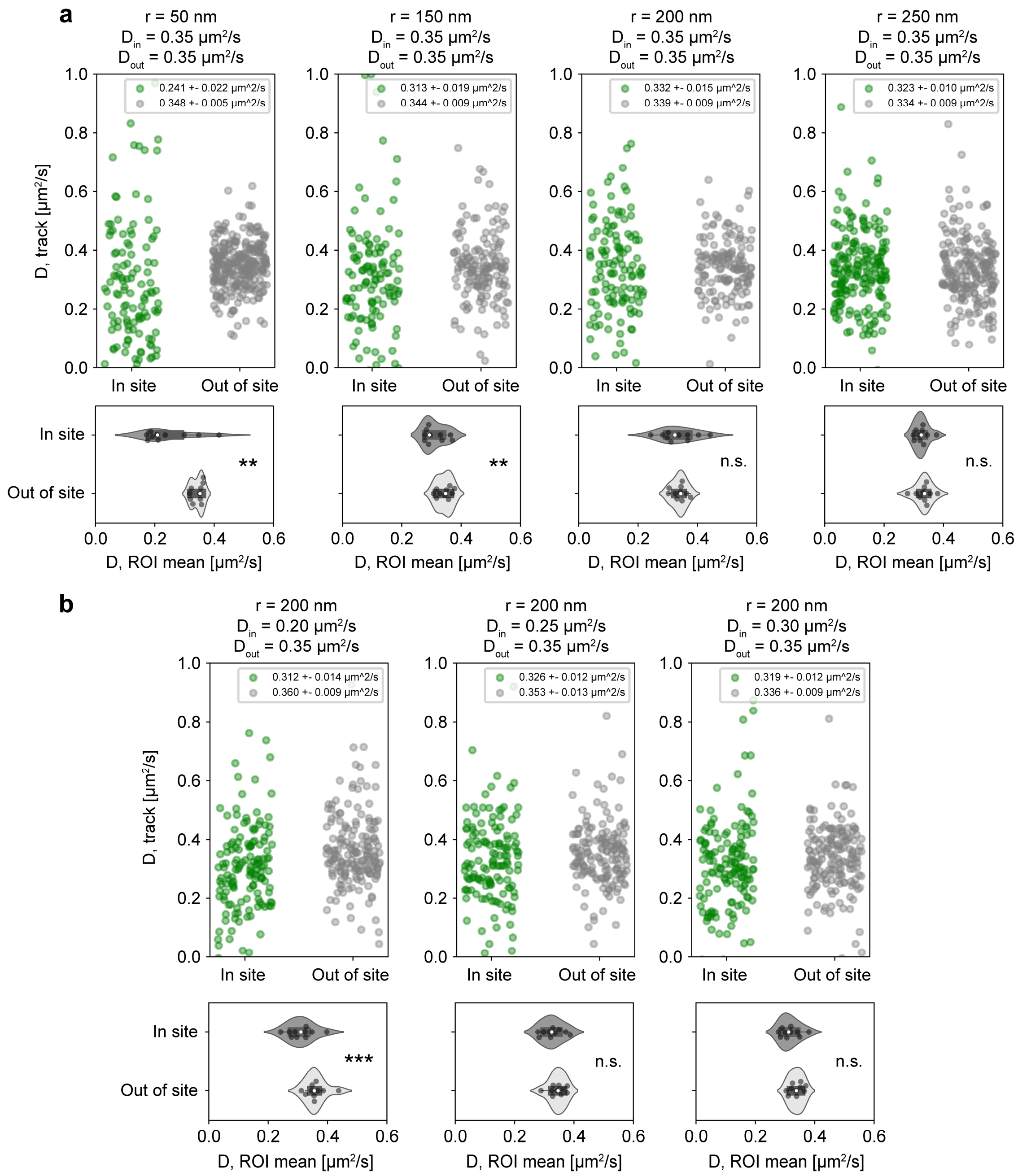


**Suppl. Figure 8. Diffusion analysis on simulated data.** Results from diffusion coefficient analysis performed on simulated data with various parameter combinations. **a.** Varying analysis site radius, from 50 nm to 250 nm, showing bias in diffusion coefficient estimation for smaller site radii. **b.** Varying simulated inner diffusion coefficient, from 0.20 to 0.35 µm$^2$/s, showing sensitivity to detecting diffusion coefficient variations for smaller values. p-values: (a) r = 50: p = 0.0060a; r = 150: p = 0.0084; r = 200: p = 0.86; r = 250: p = 0.41, (b) $D_{in}$=0.20: p = 0.00086; $D_{in}$ = 0.25: p = 0.35; $D_{in}$ = 0.30: p = 0.096. For each condition either 10 ROIs, each with 100 tracks, each with a mean of 200 localizations, or 20 ROIs, each with 250 tracks, each with a mean of 200 localizations, were simulated.

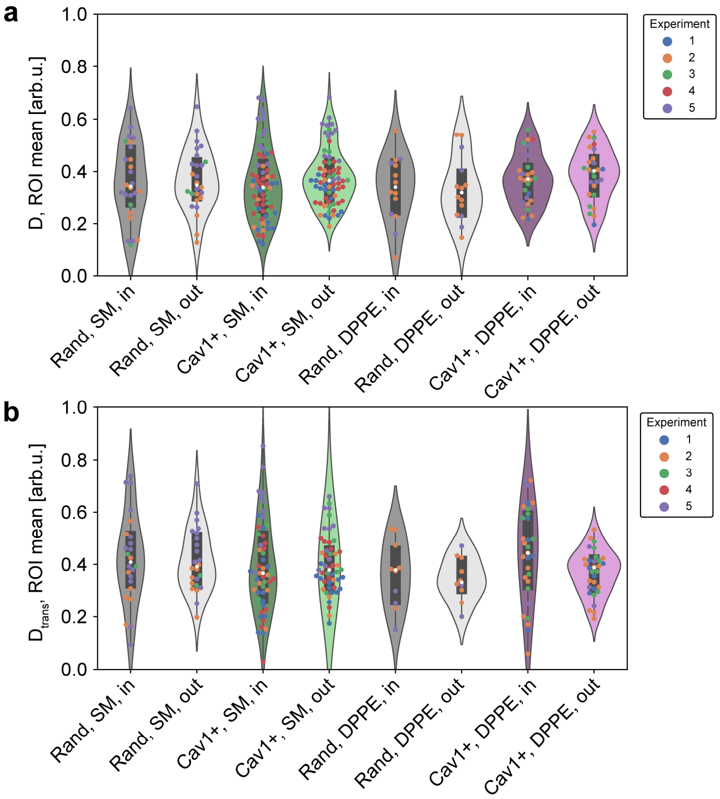


**Suppl. Figure 9. Diffusion analysis results from Caveolin1 accumulation sites.** Results from diffusion coefficient and transient diffusion coefficient analysis, showing the extracted diffusion coefficients inside and outside the site. Each datapoint represents the mean from all tracks in a ROI, and different experiments are color-coded. Groupings, from left to right, are inside and outside randomized sites for SM tracking, inside and outside Caveolin1-triggered sites for SM tracking, inside and outside randomized sites for DPPE tracking, and inside and outside Caveolin1-triggered sites for DPPE tracking. **a.** Diffusion coefficient analysis. **b.** Transient diffusion coefficient analysis.

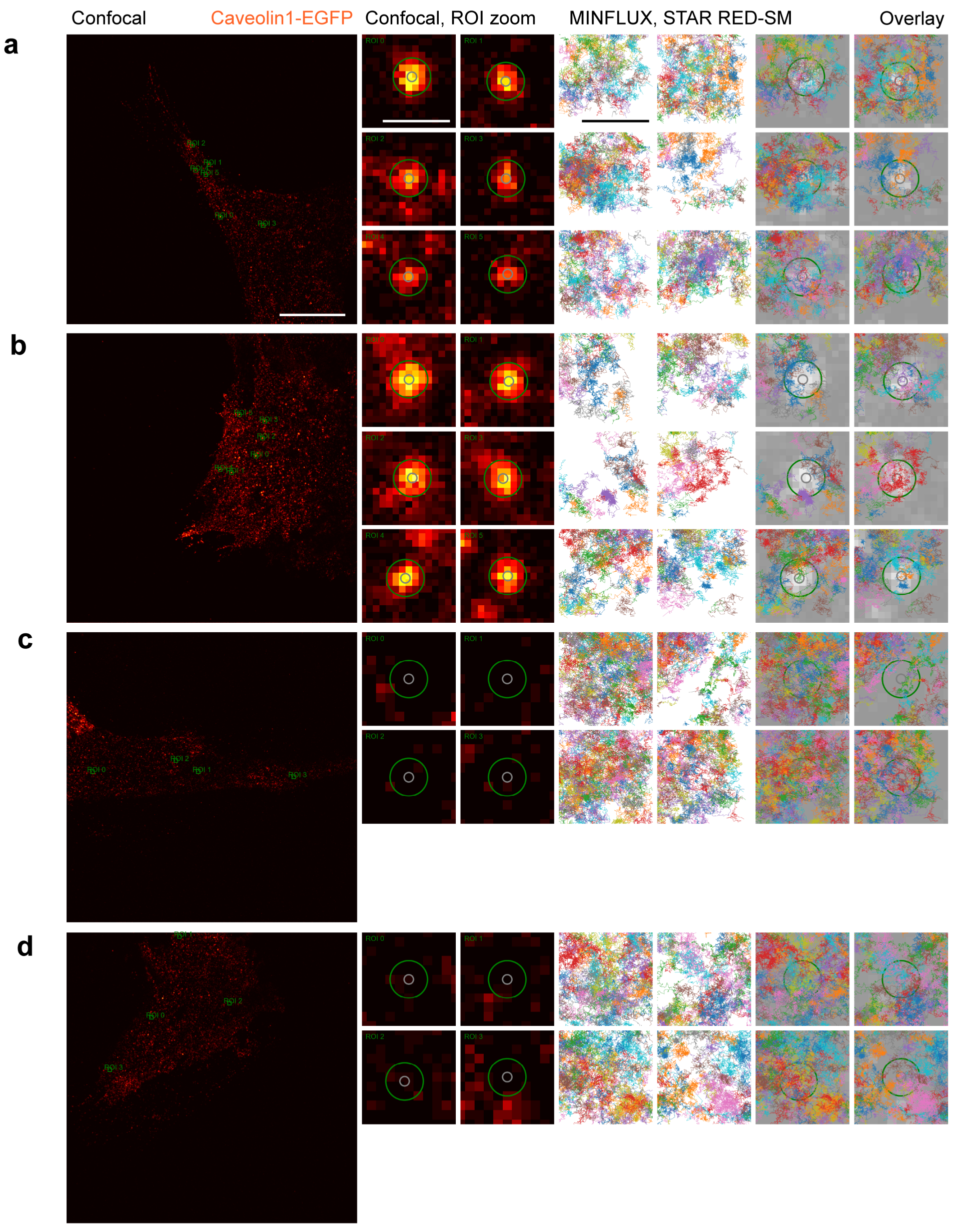


**Suppl. Figure 10. etMINFLUX caveolae and random sites with SM-STAR RED – further examples.** Further examples of (**a**,**b**) Caveolin1 event detection and (**c**,**d**) random site detection, and MINFLUX tracking of a lipid analogue (SM-STAR RED) in small ROIs around detected event or random sites. Scale bars: 5 µm (confocal overviews), 250 nm (confocal zooms; MINFLUX tracks).

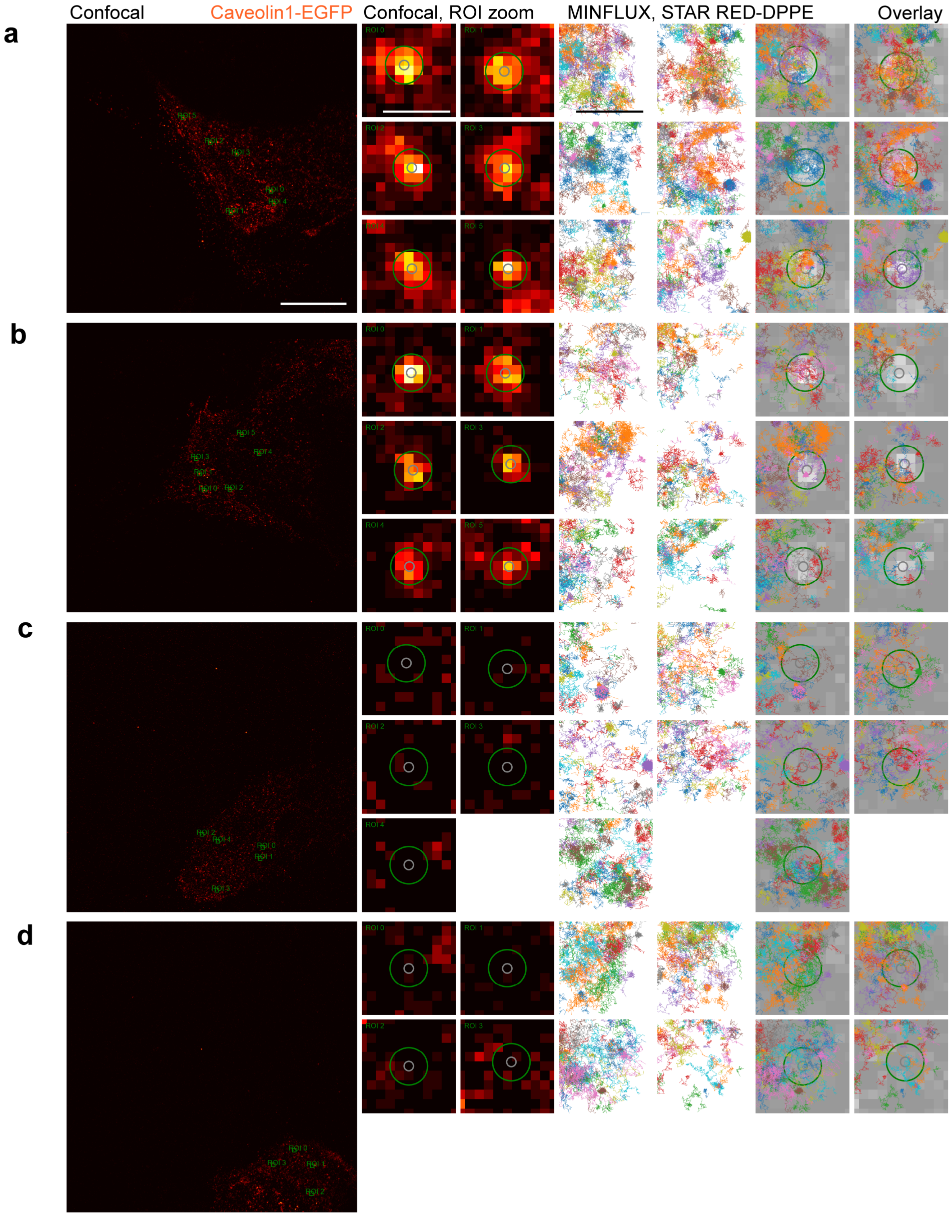


**Suppl. Figure 11. etMINFLUX caveolae and random sites with DPPE-STAR RED – further examples.** Further examples of (**a**,**b**) Caveolin1 event detection and (**b**,**d**) random site detection, and MINFLUX tracking of a lipid analogue (DPPE-STAR RED) in small ROIs around detected event or random sites. Scale bars: 5 µm (confocal overviews), 250 nm (confocal zooms; MINFLUX tracks).

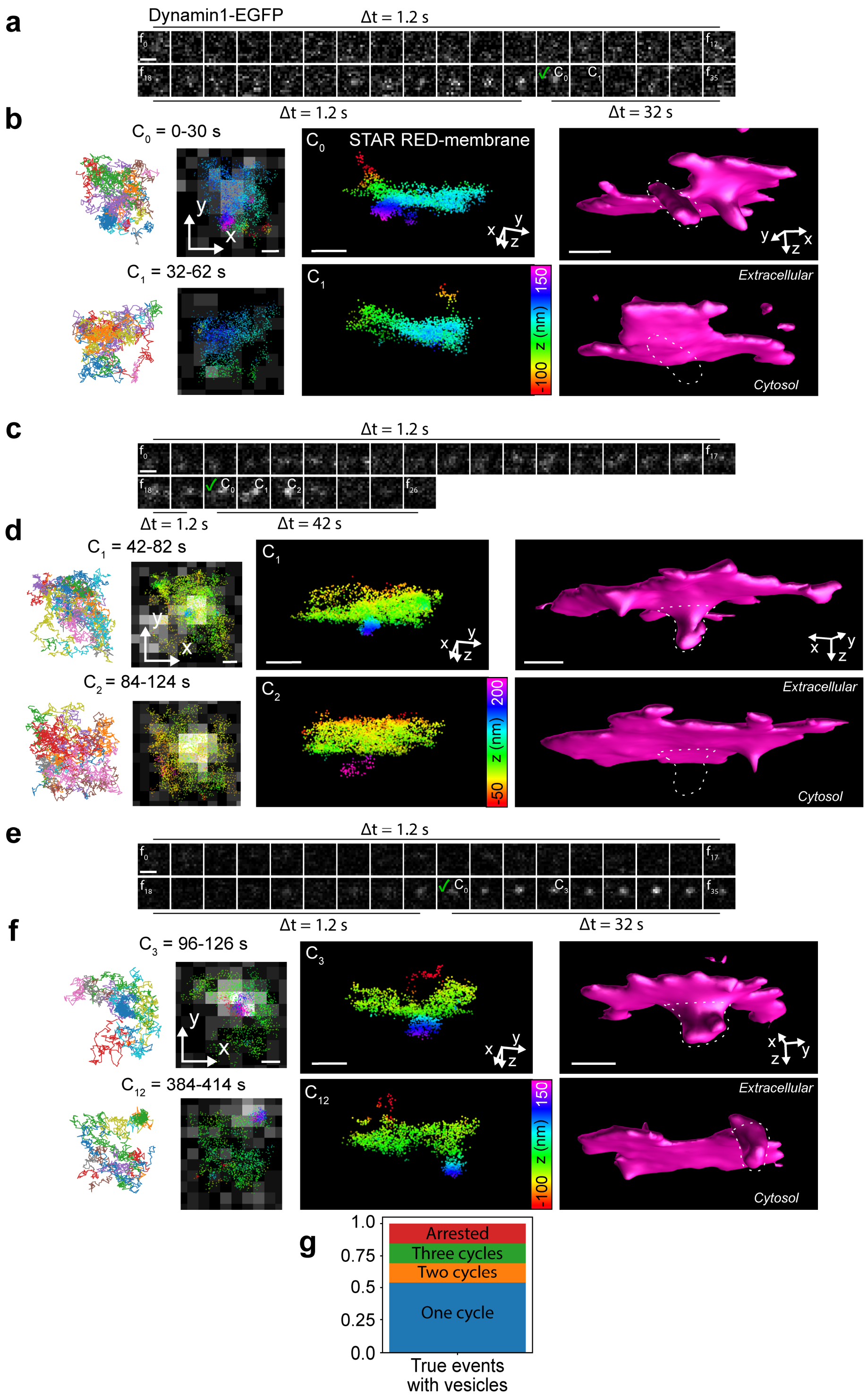


**Suppl. Figure 12. etMINFLUX endocytosis sites – further examples. a.** Further examples of Dynamin1 accumulation event detection and MINFLUX tracking of a membrane marker in small ROIs around detected event sites, with topological maps of the event site. Scale bars: 500 nm (**a, c, e**), 100 nm (**b, d, f**).

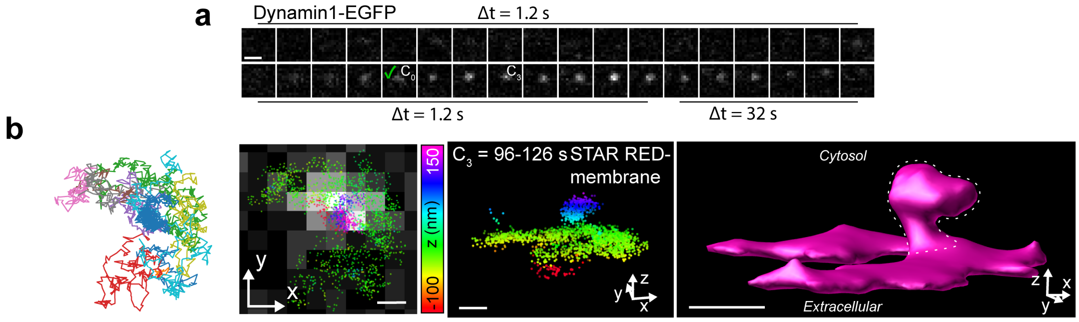


**Suppl. Figure 13. etMINFLUX endocytosis site with long endosome neck. a.** A further example of a Dynamin1 accumulation event detection and MINFLUX tracking of a membrane marker in a small ROI around the detected event site. **b.** MINFLUX tracking data from the event, represented as tracks, localization clouds in 2D and 3D, as well as a topological map of the membrane surface. Scale bars: 500 nm (**a**), 100 nm (**b**).

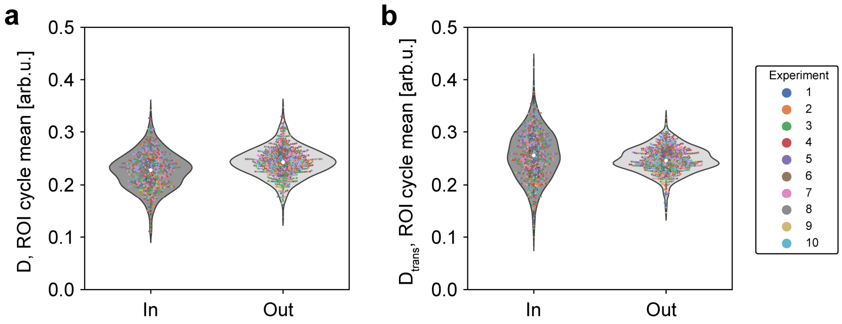


**Suppl. Figure 14. Diffusion coefficient and transient diffusion coefficient analysis results from Gag accumulation sites.** Results from diffusion coefficient and transient diffusion coefficient analysis, showing the extracted diffusion coefficients inside and outside the site. Each datapoint represents the mean from all tracks in a ROI cycle, and different experiments are color-coded. **a.** Diffusion coefficient analysis. **b.** Transient diffusion coefficient analysis.

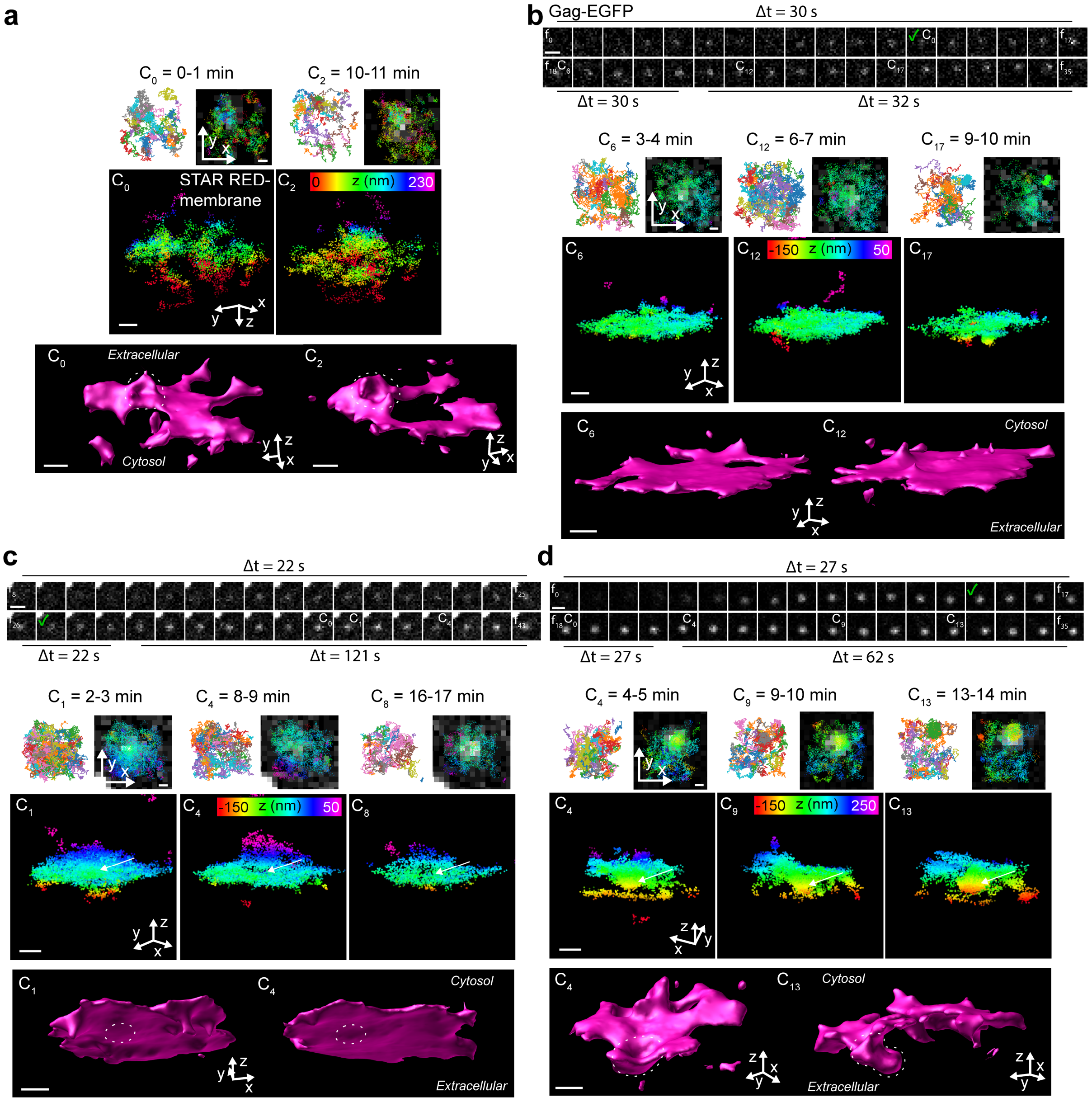


**Suppl. Figure 15. etMINFLUX virus budding sites – further examples.** Further examples of Gag accumulation event detection and MINFLUX tracking of a membrane marker in small ROIs around detected event sites, with topological maps of the event site. **a.** Additional time points of the event shown in Fig. 5f,h. **b.** An event with no sign of budding. **c.** An event with moderate static bulging. **d.** An event with a developing budding site. Scale bars: 500 nm (confocal time lapse in **b–d**), 100 nm (MINFLUX localizations-confocal overlay, 3D MINFLUX localizations, and 3D MINFLUX surface in **a–d**).

### Supplementary Tables

**Supplementary Table 1. etMINFLUX analysis pipeline parameter value ranges.**

| ***peak_detection*** | | | | | | | | | | | | | |
|---|---|---|---|---|---|---|---|---|---|---|---|---|---|
| **maxfilter _kersize** | **min_ dist** | **th_abs** | **sm_ rad** | **border _lim** | **init_sm** | **roi_ border** | **roi_th_ factor** | | | | | | |
| 5.0 | 8.0–30.0 | 2.0–10.0 | 1.0 | 15 | True | 5.0 | 6.0 | | | | | | |
| ***dyn_signalrise*** | | | | | | | | | | | | | |
| **min_dist** | **num_ peaks** | **th_abs _lo** | **th_abs _hi** | **border _lim** | **mem_f rames** | **tr_s_di st** | **frames _app** | **th_inc rat** | **th_incrat_ max** | **th_move _dist** | | | |
| 1.0 | 200–1000 | 0.4–1.3 | 10.0–15.0 | 15 | 10 | 6–8 | 4–7 | 1.17–1.3 | 5 | 1.4–1.5 | | | |
| ***gag_signalrise*** | | | | | | | | | | | | | |
| **min_dist_ app** | **num_ peaks** | **th_abs _lo** | **th_abs _hi** | **finalint _lo** | **finalint _hi** | **border _lim** | **mem_f rames** | **tr_s_d ist** | **frames_app** | **th_incrat** | **th_incrat _max** | **incslope** | **th_move _dist** |
| 1.0–5.0 | 100–500 | 1.3–5.0 | 10–300 | 0.75 | 5.0–50.0 | 10 | 5–6 | 10 | 3–6 | 1.1–1.3 | 5.0–15.0 | 0.05–0.1 | 1.3–3 |

**Supplementary Table 2. Confocal and MINFLUX acquisition parameters in the various experiments.**

| | Confocal | | | | | | MINFLUX | | | |
|---|---|---|---|---|---|---|---|---|---|---|
| **Experiment** | **Exc. power, 488 nm, sample [µW]** | **ROI size [µm$^2$]** | **Pixel size [nm]** | **Pixel dwell time [µs]** | **Frame time [s]** | **Frame period [s]** | **Exc. power, it. 1, 640 nm, sample [µW]** | **ROI size [µm$^2$]** | **Cycle time [s]** | **Cycle period [s]** |
| Caveolin1 | 1–10 | 15×15 or 20×20 or 80×80 | 70 or 100 | 2 | 0.3 or 0.4 or 1.9 or 3.2 | - | 20–60 | 1–2×1–2 | 52–90 | - |
| Dynamin1 | 0.6–3.0 | 10×10 or 15×15 | 70 | 20 | 0.6 or 1.2 | 0.6–1.2 | 39–40.5 | 0.5–0.8×0.5–0.8 | 20–40 | 21–42 |
| Gag | 0.7–2.7 | 10–25×10–25 | 60 | 10 | 0.5–2.1 | 20–30 | 39.0–39.9 | 0.7–0.8×0.7–0.8 | 30–60 | 31–298 |
